# Coexistence in diverse communities with higher-order interactions

**DOI:** 10.1101/2022.03.04.483022

**Authors:** Theo Gibbs, Simon A. Levin, Jonathan M. Levine

## Abstract

A central assumption in most ecological models is that the interactions in a community operate only between pairs of species. However, the interaction between two species may be fundamentally changed by the presence of others. Although interactions among three or more species, called higher-order interactions, have the potential to modify our theoretical understanding of coexistence, ecologists lack clear expectations for how these interactions shape community structure. Here, we analytically predict and numerically confirm how the variability and strength of higher-order interactions affect species coexistence. We found that, as higher-order interaction strengths become more variable across species, fewer species coexist, echoing the behavior of pairwise models. If inter-specific higher-order interactions become too harmful relative to self-regulation, coexistence was destabilized, but coexistence was also lost when these interactions were too weak and mutualistic effects became prevalent. Last, we showed that more species rich communities structured by higher-order interactions lose species more readily than their species poor counterparts, generalizing classic results for community stability. Our work provides needed theoretical expectation for how higher-order interactions impact species coexistence in diverse communities.

## Introduction

A fundamental problem in ecology is explaining species coexistence in diverse communities despite the force of competitive exclusion. Because of the inherent complexity of diverse systems, research on this problem has typically advanced by assuming that interactions operate only between pairs of species, and that these pairwise interactions then combine to generate the dynamics of the full community [1]. When communities have more than two species, however, any pairwise interaction can be modified by one or more of the other community members. Such interactions, termed higher-order interactions, are absent from the purely pairwise models that have contributed most to our understanding of species coexistence. Despite longstanding efforts in ecology [2, 3, 4, 5, 6, 7] and other fields [8, 9, 10, 11, 12], we currently lack coherent theoretical expectations for how higher-order interactions impact coexistence.

Higher-order interactions emerge when species plastically respond to other species in ways that affect their interaction with still other species. For example, an early-season plant species may delay the growth period of a mid-season competitor, causing the latter to compete more strongly with a late-season species [1]. Interactions of this type are presumably frequent in nature, and empirical evidence for their operation in plant and microbial communities is accumulating [13, 14, 15]. While demonstrating the operation of higher-order interactions is the key first step in this research area, the obvious follow-up question is how higher-order interactions actually shape the dynamics of the community.

Ecologists do know how interactions shape coexistence in the simplest possible case of just two competing species, and these rules form a null expectation for the influence of higher-order interactions [16]. These rules include:

1. Coexistence is favored when intra-specific competition is stronger than inter-specific competition.
2. Coexistence is disfavored by large variance among species in their intrinsic growth rates and sensitivities to competition.
3. Species abundances grow without bound when inter-specific interactions are facilitative rather than harmful, and stronger than self-regulating interactions.

Generalizing these rules to diverse ecosystems – even those with only pairwise interactions – is mathematically challenging, as the coexistence outcomes depend on the full structure of the competitive network [17]. One feasible path forward involves first ignoring this structure, and analyzing how the summary statistics of pairwise interactions affect coexistence. Taking this approach, ecologists have shown that the qualitative rules for the dynamics two competing species can still apply in diverse communities with pairwise interactions [18, 19, 20, 21, 22, 23, 24]. Understanding how the statistics of higher-order interactions influence coexistence could refine our theories of species diversity in general. Indeed, recent theoretical work studying communities of fixed total abundance [25] suggests that higher-order interactions may upend classical expectations related to diversity and stability, though whether this extends to systems where the total abundance emerges from the interactions themselves remains to be explored.

Here, we combine numerical simulation with a technique from statistical physics [21, 26, 27, 28, 29] to address three questions: (1) How do the strength and variability of higher-order interactions affect species coexistence in diverse communities? (2) How do these effects compare to the rules for coexistence in pairwise systems? (3) Does considering higher-order interactions alter classical theoretical results relating diversity to the probability of coexistence [18, 19, 20]?

## Results

We explored our research questions with a simple extension of the generalized Lotka-Volterra model to include third-order interactions [30, 31, 32], similar to those used in recent empirical studies [13, 14, 15]. In a community with *S* species, the dynamics of species *i*’s density, *N*_*i*_, is given by

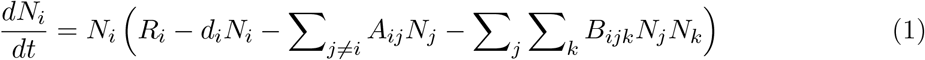

where *R*_*i*_ is species *i*’s intrinsic growth rate, *d*_*i*_ describes the strength of intra-specific limitation (set to 1 from now on for simplicity), and the coefficient *A*_*ij*_ describes the pairwise impact of species *j* on species *i*’s growth (Fig. 1(A)). We model the higher-order interactions experienced by species *i* as the product of two other species’ densities *N*_*j*_ and *N*_*k*_. *B*_*ijk*_ measures the impact of species *k* on the pairwise effect of species *j* on species *i*. We allow higher-order interactions to include squares, but not cubes, of abundances (*B*_*iii*_ = 0 for all *i*) because we assume that species’ intra-specific regulation is fully captured by the pairwise parameter *d*_*i*_, though we consider the case of cubic self-regulation later in the Results. The sign of the coefficients *A*_*ij*_ and *B*_*ijk*_ can be either positive (and therefore harmful) or negative (and mutualistic).

**Figure 1:**
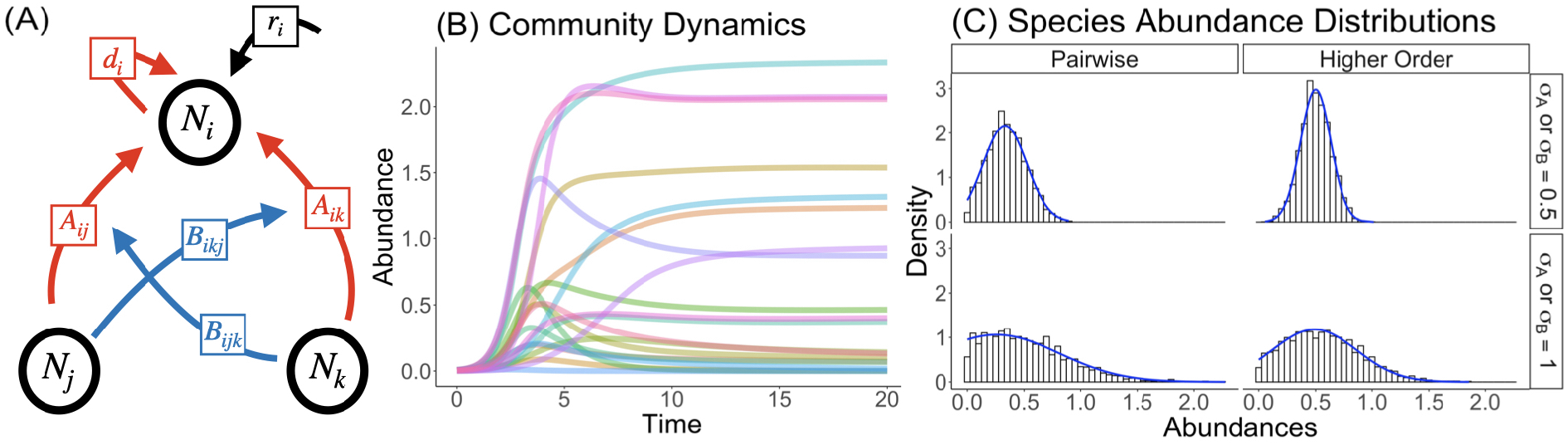
(A) Following Equation 1, each species *i* in the community has an intrinsic growth rate *R*_*i*_, experiences self-limitation through the parameter *d*_*i*_ and competes with other species through pairwise (red) and higher-order (blue) interactions. The pairwise competition coefficient *A*_*ij*_ measures the impact of species *j* on species *i*, while the higher-order coefficient *B*_*ijk*_ measures how species *k* modifies the effect of species *j* on species *i*. (B) In our simulations, we integrate the dynamics of Equation 1 with either or both pairwise and higher-order interactions, and record the fraction of species that are excluded and the abundances of the coexisting species at equilibrium. (C) Plots of the density of species with a given abundance (the species abundance distribution) for communities with either pairwise or higher-order interactions, with different interaction variances. The solid blue curves denote the predicted species abundance distributions from the cavity method (see the Methods for details).

Because of the complexity of ecological models with large numbers of species, we aim to understand how species coexistence depends on summary statistics of the species’ growth rates and interactions, rather than any particular parameterization. To understand how the mean and variance of the growth rates (*µ*_*R*_ and 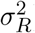) and pairwise (*µ*_*A*_ and 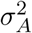) and higher-order (*µ*_*B*_ and 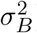) interactions influence coexistence, we varied these statistics and used both numerical simulations and analytical theory to determine how they influence the fraction of coexisting species. Because there are many more higher-order interactions than pairwise interactions among the *S* species, we scale the interaction strengths to account for the number of pairwise or higher-order interactions (see the Supplementary Information for a more complete discussion). In our simulations, we drew the growth rates and interactions from normal distributions with the specified means and variances, solved the dynamical system in Eq. 1 numerically and then recorded the abundances of all species in the community (Fig. 1(B)). This was then repeated many times for different means and variances to understand how these factors influence species coexistence. We concentrate on systems whose interactions are on average harmful but, due to their variance, can sometimes be mutualistic. Here, rather than asking whether the *S* species equilibrium is stable or feasible, as in most previous work [18, 19, 20, 22, 23, 24, 33, 34], we ask what fraction of the species coexist (*ϕ*) after the dynamics proceed from positive, randomly assigned initial species densities.

In addition to the simulation results, we use the cavity method, a technique from statistical physics [26, 27], to generate analytical predictions for how species coexistence depends on the mean and variance of species’ growth rates, pairwise and higher-order interactions. In recent work, the cavity method has successfully predicted the equilibrium species abundance distributions for a wide variety of ecological models [21, 28, 29, 35, 36, 37, 38, 39]. This method is only theoretically valid in the limit of an infinite number of species (*S* → ∞), but it has been shown to describe the behavior of relatively small Lotka-Volterra communities with pairwise interactions [21, 29]. We use the cavity method to calculate the species abundance distribution at equilibrium for a range of interaction statistics. The fraction of species with positive abundances then yields the fraction of coexisting species *ϕ* (see the Methods for the equations and the Supplementary Information for the full calculation). Importantly, our predicted abundance distributions closely match their simulated counterparts (Fig. 1(C)).

Our first main result is that the cavity method does an excellent job predicting the fraction of coexisting species for the Lotka-Volterra model with higher-order interactions. When the community was very small (only 5 species), we found considerable divergence between the simulation outcomes and the cavity method predictions, and thus our theory is not predictive for such small systems (Fig. 2). At the other extreme, when the system began with 30 species, or with much larger species pools (*S* = 300) as shown in the Supplementary Information, we found excellent agreement between our simulations and the cavity method predictions (Fig. 2). Importantly, even when there were only 10 species, the cavity method captured the average behavior of the simulated communities.

**Figure 2:**
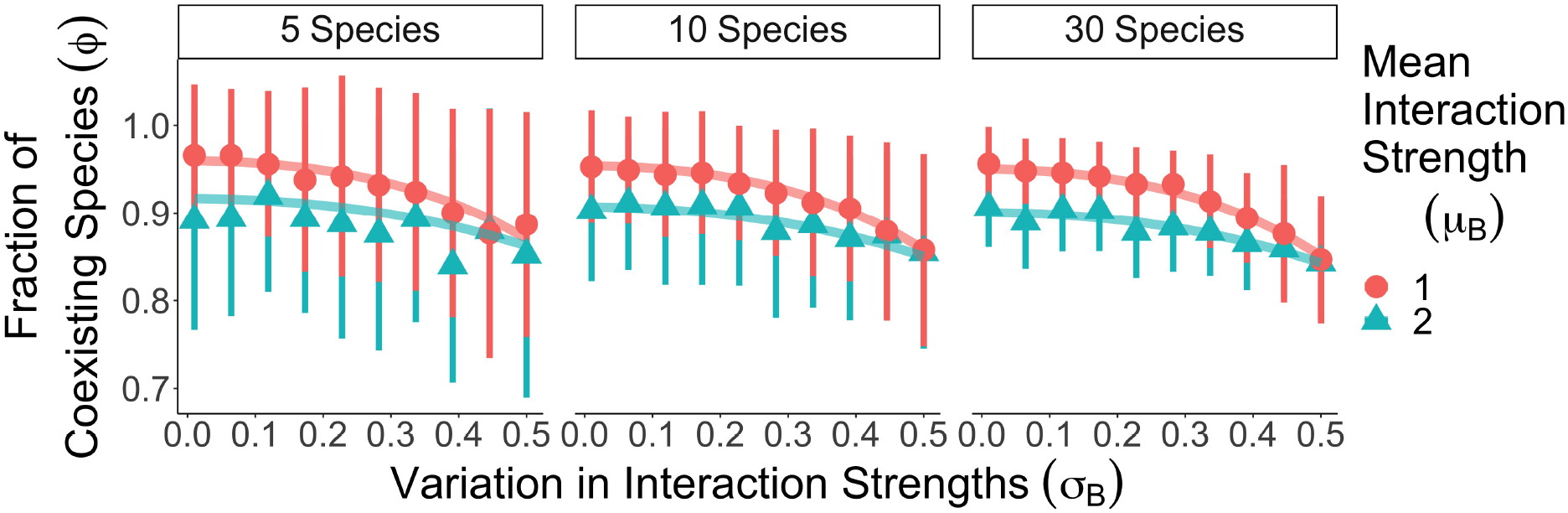
Comparison between simulated fraction of coexisting species, shown by symbols +/- 1 standard deviation, and cavity method predictions of that fraction, shown by the continuous lines. Results are shown for different community sizes, and mean and variance of higher-order interaction strength. Simulation means and standard deviations were obtained for 100 initial communities per parameter combination. In all panels, the mean intrinsic growth rate was *µ*_*R*_ = 1.5 and its variance was *σ*_*R*_ = 0.5.

We first explored the effects of the pairwise interaction strengths and variances on species coexistence to provide a baseline against which to compare the effects of higher-order interactions. In pairwise systems, increasing the variance in the interaction strength, reduces the fraction of species that coexist. This happens because, just like in models with only two competitors, variability in the inter-specific interaction strengths favors some competitors over others (some get a better draw of interactions than others), driving the losers to exclusion (Fig. 3(A) and [21, 28, 29]). Increasing the variation in growth rates has an analogous effect, which can be seen, for example, by comparing the blue lines (where mean interaction strength is fixed) across the top three panels of (Fig. 3(A) and [21, 28, 29]).

**Figure 3:**
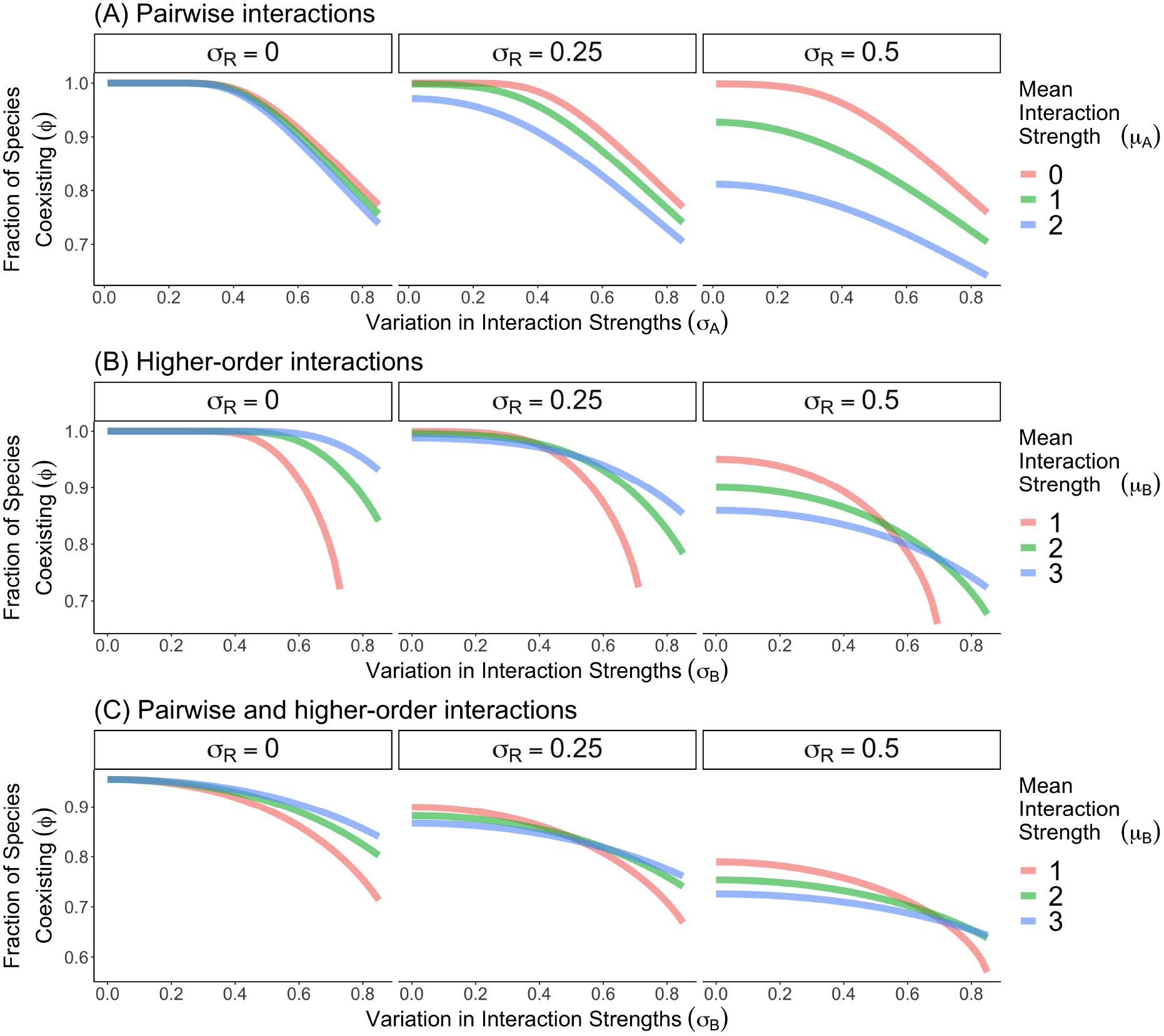
Predictions for how the fraction of coexisting species (*ϕ*) depends on the variation in interaction strengths assuming only pairwise (Panels A), only higher-order (Panels B) or both types of (Panels C) interaction in the system. We plot results for three different variation in intrinsic growth rates (*σ*_*R*_) and three different mean interaction strengths (*µ*_*A*_ or *µ*_*B*_). In Panels C, the pairwise interaction statistics are *µ*_*A*_ = 1 and *σ*_*A*_ = 0.5. In all panels, the mean growth rate is *µ*_*R*_ = 1.5. Simulation results in the Supplementary Information closely match these predictions. Throughout our work, we implement a correction to the standard cavity method equations so that we are better able to predict coexistence in smaller communities (see Methods and Supplementary Information).

Increasing the mean pairwise interaction strength, while keeping the intra-specific interaction strength constant (*d*_*i*_ = 1) reduces coexistence because the inter-specific interactions become on the whole more competitive (Fig. 3(A) and [21, 28, 29]). With more species exerting greater effects on others than on themselves, the system exhibits less coexistence. This decrease in coexistence becomes most apparent when species’ competitive imbalances increase, either through variation in the intrinsic growth rates or variation in interaction strengths. We can attribute this behavior to the relative strength of intra- to inter-specific interactions because, if instead species with a common intrinsic growth rate experience a concomitant increase in self-regulation as their inter-specific interactions become more competitive, the mean competition strength has no impact on coexistence (see Supplementary Information and [21, 28]).

Our second main result is that some of the lessons from how pairwise interactions affect coexistence translate over to the effects of higher-order interactions. For example, increased variability in higher-order interaction strength (and intrinsic growth rates) reduced species coexistence (Fig. 3(B)) because, just as in the pairwise case, this variability favors some species over others, and the losers get excluded. Similarly, as long as the variation in higher-order interaction strength was relatively low, and species differed in their intrinsic growth rates, making higher-order interactions more harmful on average while keeping self-regulation constant reduced coexistence.

However, our third main result is that higher-order interaction strength differed markedly from pairwise interaction strength in its effects on species coexistence in one important way. Note that when species shared identical intrinsic growth rates or variation in the higher-order interaction strength was high, more harmful higher-order interactions increased, rather than decreased coexistence (Fig. 3(B)). This coexistence-promoting effect of more harmful higherorder interactions contrasts with the effect of more harmful pairwise interactions, and extends to systems with a mix of pairwise and higher-order interactions (Fig. 3(B-C)).

The beneficial effect of more harmful higher-order interactions emerges because such interactions reduce the likelihood of a mutualism that can cause the abundances of some species to grow without bounds. When higher-order interactions are only weakly harmful on average but highly variable, some species experience net facilitation from the most abundant species, creating runaway abundances when the mutualisms are reciprocal. This is analogous to what can emerge in two-species systems when the inter-specific mutualisms are stronger than the self-regulating terms. Indeed, the dominant species in communities with only weakly harmful higher-order interactions tended to be those that facilitate one another, while differentially harming the species with lower abundances (see Supplementary Information). This effect occurs for both pairwise and higher-order interactions (see [40, 41] and Supplementary Information), but because higher-order interactions scale with the square of the average abundance, the effect of strong inter-specific mutualism becomes more pronounced. In fact, when higher-order interactions are on average mutualistic and self-regulation is relatively weak, the mean abundance grows indefinitely. In sum, even though more harmful inter-specific interactions should decrease coexistence by overpowering intra-specific regulation, they simultaneously decrease the likelihood of runaway mutualisms, which greatly benefits coexistence.

To further evaluate this hypothesis, we simulated a model where higher-order interactions less easily generate runaway abundance because they saturate with species density. Specifically, the scalar *B*_*ijk*_ of Eq. 1 was replaced by 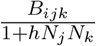 so that higher-order interactions saturate at a rate controlled by the parameter *h*. If *h* = 0, we recover the model in Eq. 1, while if *h >* 0, the higher-order interaction strengths level off with increasing densities. With this modification, the higher-order interaction properties affect coexistence in a manner identical to the pairwise interactions. Namely, more harmful higher-order interactions simply lead to less coexistence (Fig. 4). In short, modeling higher-order interactions with saturating functional responses eliminated the counter-intuitive effect of mean higher-order interaction strength, while preserving the effects of variation in the growth rates and interactions. In the Supplementary Information, we show that introducing cubic self-regulation has a similar effect on coexistence as saturating higher-order interactions.

**Figure 4:**
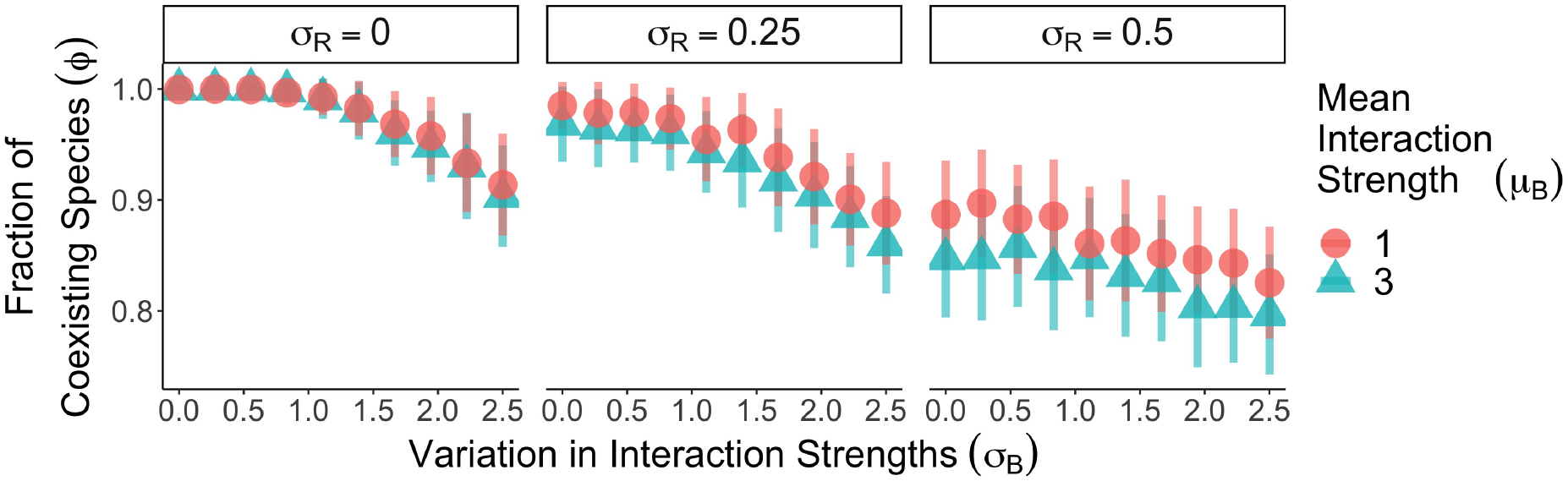
Plotted are the fraction of coexisting species for three variations in intrinsic growth rate (*σ*_*R*_) when the level of saturation is *h* = 3. The symbols show the mean of the simulation results for over 100 replicate communities and the error bars denote one standard deviation. In all panels, the number of species *S* = 30, the mean growth rate *µ*_*R*_ = 1.5, the mean pairwise interaction strength *µ*_*A*_ = 1 and the variation in pairwise interaction strengths *σ*_*A*_ = 0.25.

Our final result is that the classic finding that the probability of coexistence of all *S* species declines with species richness in systems with pairwise interactions [18, 19, 20, 42] also holds in systems with higher-order interactions. Theory predicts that, as the number of species increases, communities can tolerate less variability in their interactions before losing their first species [19, 22, 24]. We call the value of the interaction variability at which the first species goes extinct the “critical interaction variability,” denoted 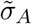 and 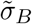 for pairwise and higher-order interactions respectively. Notably, recent theory found that under three-way higher-order interactions, this critical interaction variability exhibited no systematic dependence on the number of species [25], albeit using a different mathematical framework from the one we have considered. By contrast, in simulations of our model, the critical interaction variability for both pairwise and higherorder interactions decreased as a function of the number of species in the community (Fig. 5). In fact, the critical higher-order interaction variability decreased more quickly with species richness than the pairwise variability. Note that to obtain this result, we removed the scalings on the interaction statistics thus far imposed to remove any effect of diversity on the total effect of the interactions. Predictions for the critical variability from our cavity method framework closely matched the simulation results (see the lines in Fig. 5 and the Methods section for the corresponding equations).

**Figure 5:**
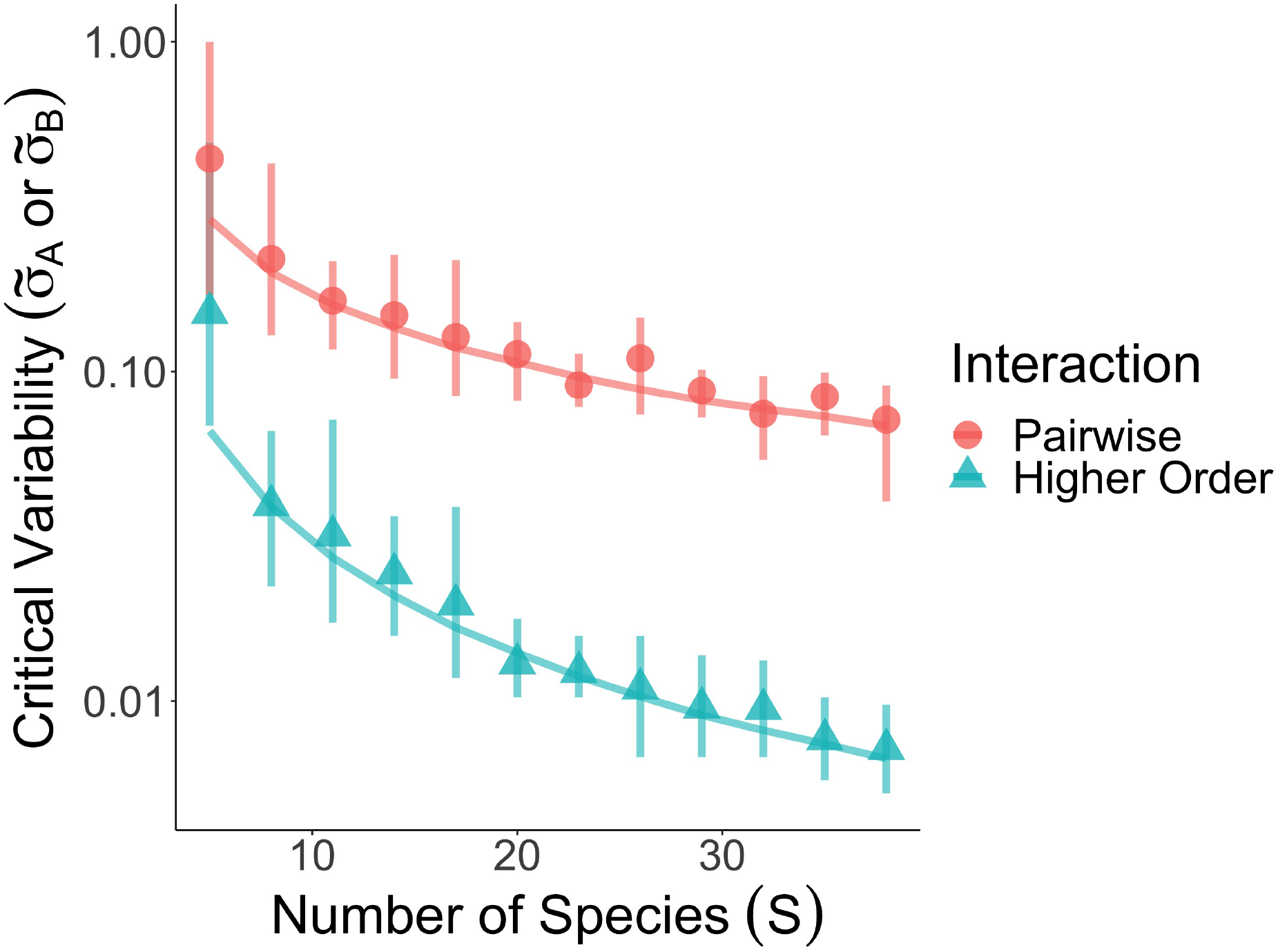
The critical interaction variability (the smallest value of 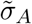 or 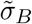 at which the first species goes extinct) as a function of the number of species in a community with either pairwise or higher-order interactions. The circles and triangles show the average of simulation results for 10 replicate communities and the error bars denote the minimum and maximum value found in all 10 simulations. The lines are predictions based on our cavity method framework in Eq. 4. We set *σ*_*R*_ = *µ*_*A*_ = *µ*_*B*_ = 0 to make the comparison with previous theory more direct and we set the mean growth rate to be *µ*_*R*_ = 1.5.

## Discussion

Through the analysis of models for two competing species, ecologists have derived simple rules for species coexistence (spelled out in the Introduction) [16, 43]. We know that these rules do not formally apply to systems with more than two species, including those with purely pairwise interactions [17]. Nonetheless, our findings here suggest that these simple rules may strongly guide expectations for the interpretation of coexistence in large systems, even those organized by higher-order interactions, as long as the network of interactions has a random structure. Moreover, the cavity method can be used to develop theory for how higher-order interactions impact species coexistence in such systems.

We found that as higher-order interaction strengths become more variable between the species, fewer species coexisted. When interactions are heterogeneous and randomly assigned to species, species differ in their sensitivity to competition, and the poorest competitors get excluded. This behavior is directly analogous to results from pairwise interactions [18, 19, 20, 21, 22]. The average strength of higher-order interactions, on the other hand, exhibited a more complex effect on coexistence. When species differed considerably in their intrinsic growth rates, and interaction strengths had little variation, more harmful inter-specific higher-order interactions generated less coexistence. This result follows from the two species rule that more harmful inter-specific interactions relative to intra-specific regulation are destabilizing [21, 22].

However, when the variation in higher-order interactions was large relative to the variation in intrinsic growth rates, we found the opposite dependence – more harmful higher-order interactions counter-intuitively produced more coexistence. Even here though, the two-species rules prove useful. Less harmful mean higher-order interactions introduced more mutualistic interactions, which even in two species systems can cause abundances to grow without bound when they overwhelm self-regulation. The subset of species engaged in this mutualistic rise were the most abundant (see Supplementary Information), and greatly harmed species that happened to engage in harmful higher-order interactions with these species. When the strength of higher-order interactions instead saturated as a function of species abundance, more harmful higher-order interactions once again produced less coexistence, because the possibility of strong mutualistic interactions, and hence groups of highly dominant species, was reduced.

In sum, while variation in higher-order interactions harmed coexistence regardless of whether or not these interactions saturated with species densities, the effect of the mean interaction strength depended strongly on the model form. An important direction for future work, therefore, is to understand how the effect of the higher-order interaction strength depends on competitor densities in the field, since this dependence strongly determines the effect of the measured higher-order interaction strengths. More generally, we suspect that the unbounded higher-order interactions currently being fit to data due to very reasonable data limitations [13] would generate unrealistic community dynamics.

Our message thus far – that the rules for coexistence under pairwise interactions apply to large communities with higher-order interactions – also holds when exploring how species richness affects opportunities for coexistence. In contrast to classic results that communities tolerate less variability in pairwise inter-specific interactions as they become more diverse [18, 19, 20], Bairey *et al*. [25] recently showed that species diversity has no effect on the variability of three-way, higher-order interactions that disrupt coexistence. In our model, the critical variability of both pairwise and higher-order interactions actually declined with species richness. In fact, the critical higher interaction variability decreased more rapidly with species richness than did the critical pairwise interaction variability. We believe the discrepancy between our results and those of Bairey *et al*. [25] lies in the different modeling frameworks. In the replicator equation used by Bairey *et al*. [25], all abundances must sum to one, and thus higher-order interactions become weaker as the number of species increases because the products of relative abundances near zero quickly become very small. In the generalization of the Lotka-Volterra model we consider, every species has an abundance that is fixed by its intrinsic growth rate and its self-regulation. As a result, the variability in higher-order interactions and species richness affects opportunities for coexistence in the same qualitative way that pairwise interactions do.

More generally, our results suggest that the number of interactions in a community plays an important role in determining their effects on coexistence. When we removed the scalings which accounted for the larger number of higher-order than pairwise interactions, we found that higher-order interactions had a stronger impact on coexistence simply because there were more of them. This fact suggests that higher-order interactions involving more than three species should have even smaller critical variabilities than those we predicted for three-way higher-order interactions. On the other hand, if in nature the measured strength of higher-order interactions tends to decreases as a function of the number of species involved, then our theory that scales out the number of interactions may be a more accurate representation of real systems. The relationship between the order of an interaction and its empirically derived strength is therefore an important outstanding question for both theoreticians deriving higher-order interactions from mechanistic underpinnings and experimentalists tackling the problem in nature.

Our theory generates a null expectation for how higher-order interactions influence species coexistence assuming they are on average harmful (mutualistic, non-saturating higher-order interactions simply cause abundances to explode), and there is no structure to the higher-order interaction network. Indeed, we have shown with the cavity method that, when higher-order interactions are sampled at random, they do not generate ecosystems with perfect coexistence. However, ecological interactions in nature are likely to be non-random. If the network of higherorder interactions has some complex structure, then it may have a fundamentally different effect on coexistence than suggested here. For example, we assumed that higher-order interactions involving the square of abundances (the intra-specific higher-order interactions [13]) follow the same distribution as all other higher-order interaction terms. If instead the intra-specific higherorder interactions are stronger than their inter-specific counterparts, they might be broadly stabilizing. Similarly, higher-order interaction strength may be correlated with the underlying pairwise interactions in the system, and thereby give rise to more or less coexisting species than predicted here. At this point, however, it is unclear how to impose additional constraints on the parameters of the model we consider without specifying a mechanism for the interactions in the ecosystem [44, 45]. Moreover, deviations from truly random interactions may not alter the qualitative conclusions we have focused on here. If specific interaction structures were found to change our main conclusions, it is possible to incorporate these structures into the cavity method [28, 41], allowing one to understand the mechanisms by which non-random interaction structures benefit or harm coexistence.

In this context, a central challenge in this field is to derive phenomenological higher-order interaction parameters either from (1) nature, or (2) an underlying mechanistic process in a model. Although both approaches could refine the conclusions we have derived based on randomly-sampled interactions, an empirically-determined interaction network can be used to interrogate specific patterns in the structure of the interactions. At the same time, the number of possible higher-order interactions grows quickly with the number of species and the order of the interactions themselves, making it very difficult to estimate all possible interactions experimentally and necessitating new empirical approaches to circumvent this challenge. This is where the cavity method may prove particularly useful in an empirical context. To predict coexistence with the cavity method, one only needs estimates of the mean and variability of the interactions [41]. In other words, not every interaction needs to be measured. As a result, the cavity method provides a powerful framework for empiricists to compare the potential effects of higher-order interactions across different ecosystems. The theory can used to generate a baseline level of coexistence expected from randomly assigned higher-order interactions, and thus deviations from such predictions can be indicators of more complex ecological structure in nature.

## Methods

### Simulation details

In our simulations, we used the LSODA solver from the deSolve v1.25 package [46] in R version 3.6.1. We sample the species’ growth rates and interaction parameters from normal distributions with the statistics we specified in the Results. We start each simulation with all species present at randomly selected abundances in the interval [0, 1]. We integrate the dynamics for 10^7^ timesteps and then record the abundances. We designate species with abundance smaller than 10^*−*14^ to be extinct. We then test whether or not we have reached equilibrium by computing each of the coexisting species per-capita growth rates and comparing them to a cutoff of 0.01. We also test if any of the excluded species can invade the equilibrium using the same growth rate threshold. If the dynamics did not reach equilibrium or an excluded species can invade, we remove the simulation run for our data. This occurs rarely and only for large values of *σ*_*A*_ or *σ*_*B*_ (see Supplementary Information for further discussion of our simulation methods). In the Supplementary Information, we also test whether or not there are multiple stable equilibria in the dynamics of Eq. 1 by finding the equilibria of the model with the same parameters for a suite of different random initial conditions, and we find that this also occurs rarely and only when *σ*_*A*_ or *σ*_*B*_ is large, as in [21, 28]. The code used to run simulations and generate figures is available on GitHub at https://github.com/tgibbs-hub/CavityHOIs.

### Cavity method equations

In the Supplementary Information, we provide the cavity method calculation in detail. In this and the following sections, we discuss the calculation without the finite-size corrections that we mentioned in the main text, but see the Supplementary Information for the complete analysis. The predicted distribution for the coexisting species is a truncated normal distribution (Fig. 1(C)). We use *µ*_0_ and *σ*_0_ to denote the mean and standard deviation of this distribution before truncating it. We derive equations in the Supplementary Information for how these statistics relate to the growth rate and interaction statistics. Specifically, we find that

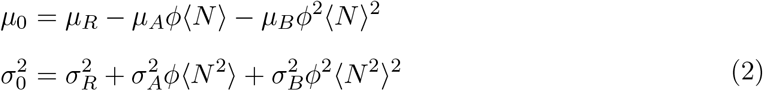

where *ϕ* is the fraction of coexisting species (as in the Results), ⟨*N* ⟩ is the mean of the coexisting species, ⟨*N* ^2^⟩ is the second moment of the coexisting species. In the Supplementary Information, we actually treat a more general case in which the pairwise interaction coefficients can be correlated across the diagonal (ie. 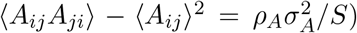), but we report the simpler formulas in which *ρ*_*A*_ = 0 here. Eq. 2 can be interpreted as the average and variance in abundances of a given coexisting species. Specifically, we can solve Eq. 1 for a focal abundance *N*_*i*_, and then compute the mean and variance of the resulting solution using both the unknown properties of the coexisting species (*ϕ*, ⟨*N* ⟩ and ⟨*N* ^2^⟩) as well as the statistics of the growth rates and interactions. In fact, previous work [47, 48] used this method to predict the equilibrium properties of the Lotka-Volterra model without invoking the cavity method. In Equation 2, *ϕ*, ⟨*N* ⟩ and ⟨*N* ^2^⟩ are all unknowns, but they are related to *µ*_0_ and *σ*_0_. Let *P* (*N*_0_| *µ*_0_, *σ*_0_) be the non-truncated normal distribution with mean *µ*_0_ and standard deviation *σ*_0_. Then, *ϕ* is the integral of *P* (*N*_0_| *µ*_0_, *σ*_0_) over the positive abundances. Similarly, ⟨*N* ⟩ is the average of *P* (*N*_0_| *µ*_0_, *σ*_0_) over the positive abundances. We find that

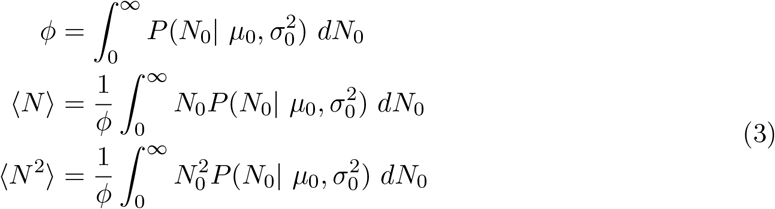

where the factors of 1/*ϕ* normalizes the integral. All in all, we have three equations for three unknowns that we can solve numerically to determine the species abundance distribution.

### The limit in which all species coexist

The formulas in Eq. 2 cannot easily be solved because *µ*_0_ and 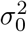 are not the same as ⟨*N* ⟩ and ⟨*N* ^2^⟩ − ⟨*N* ⟩^2^ respectively, because the values of ⟨*N* ⟩ and ⟨*N* ^2^⟩ − ⟨*N* ⟩^2^ both depend on the fraction of species that are excluded. However, when *ϕ* = 1, *µ*_0_ = ⟨*N* ⟩ and 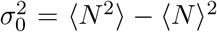 and the equations in Eq. 2 become easy to solve. Because our theory is only justified in the *S* → ∞ limit, this calculation is not fully consistent with the cavity method, since when *S* is large enough a non-zero fraction of species will always be excluded. Nonetheless, we find that it still captures the qualitative dependence of fraction of coexisting species on the mean and variance in species abundances reasonably well. In fact, in the next section, we use this limit to quantitatively predict the results of our simulation results for the critical variabilities.

When every species has the same growth rate (*σ*_*R*_ = 0) and there are only pairwise interactions in the ecosystem *µ*_*B*_ = *σ*_*B*_ = 0, we find that 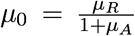 and 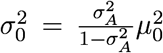. As a result, we find that *σ*_0_ is directly proportional to *µ*_0_. Because the ratio *µ*_0_*/σ*_0_ determines *ϕ* (see Supplementary Information), this calculation suggests that, at least in the limit where all species coexist, *µ*_*A*_ should have no impact on coexistence (up to finite-size corrections, which we are neglecting here). Interestingly, this dependence is true throughout the full range of *σ*_*A*_ values we consider, rather than just for small *σ*_*A*_ values when we are close to the *ϕ* = 1 limit. A straightforward calculation (which we include in the Supplementary Information) shows that the same analysis predicts the effect of mean pairwise competition strength when species do not have identical growth rates (*σ*_*R*_ *>* 0). When there are higher-order interactions (when *µ*_*B*_ and *σ*_*B*_ are non-zero), we must now solve quadratics in Eq. 2 to get *µ*_0_ and 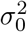. In the Supplementary Information, we find the behavior that we report in the Results – namely, that the qualitative effect of the mean higher-order interaction strength changes for different values of *σ*_0_, which is in turn determined by *σ*_*R*_ and *σ*_*B*_. Moreover, as *µ*_0_ becomes smaller, the ratio *µ*_0_*/σ*_0_ actually increases because *σ*_0_ depends on *µ*_0_. At the same time, when *σ*_*R*_ *>* 0, this dependence is reversed, and the ratio *µ*_0_*/σ*_0_ increases as *µ*_0_ increases.

### Deriving the critical variabilities

To find the critical variabilities for pairwise and higher-order interactions, we once again consider the limit where *ϕ* = 1. We use our solutions for *µ*_0_ and *σ*_0_ to compute the expected minimum of *S* samples from the normal distribution with mean *µ*_0_ and standard deviation *σ*_0_. Let *κ*(*S*) be the expected maximum value of *S* samples from the standard normal distribution. Then, the expected minimum of our predicted normal distribution is *µ*_0_ − *κ*(*S*)*σ*_0_. In our formulas, we also include the approximation 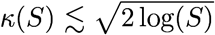, so that we can interpret the functional behavior more easily, but we use computationally determined estimates of *κ*(*S*) in Fig. 5 because they are significantly more accurate. By setting the expected minimum to zero and solving for 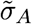 or 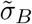, we find that the (average) critical variabilities are given by

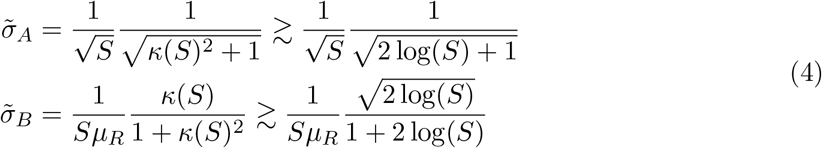

where the first factors of *S*^*−*1*/*2^ and *S*^*−*1^ respectively come from the interaction scalings that we removed in this analysis. These predictions work well (see Fig. 5), even though they are based on a series of approximations. Note that the prediction for 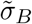 involves the mean growth rate *µ*_*R*_ because *σ*_0_ depends on *µ*_0_ which in turn depends on *µ*_*R*_, as we discussed in the previous section.

## Supplementary Information

### Supplementary Information

We provide the cavity method calculation in detail and additional simulation results.

## Acknowledgements

We thank members of the Levin and Levine labs for helpful comments and discussion. T.G. was supported by the National Science Foundation Graduate Research Fellowship Program under Grant No. DGE-2039656. Any opinions, findings, and conclusions or recommendations expressed in this material are those of the author(s) and do not necessarily reflect the views of the National Science Foundation. T.G. and J.M.L. acknowledge support from NSF Grant DEB-2022213. S.A.L. acknowledges support from NSF Grant DMS-1951358.

## References

[1] Levine, J. M., Bascompte, J., Adler, P. B. & Allesina, S. Beyond pairwise mechanisms of species coexistence in complex communities. Nature 546, 56–64 (2017). URL https://www.nature.com/articles/nature22898. Bandiera_abtest: a Cg_type: Nature Research Journals Number: 7656 Primary_atype: Reviews Publisher: Nature Publishing Group Subject_term: Ecology Subject_term_id: ecology.

[2] Abrams, P. Are Competition Coefficients Constant? Inductive Versus Deductive Approaches. The American Naturalist 116, 730–735 (1980). URL https://www.journals.uchicago.edu/doi/10.1086/283664.

[3] Adler, F. R. & Morris, W. F. A General Test for Interaction Modification. Ecology 75, 1552–1559 (1994). URL https://www.jstor.org/stable/1939616. Publisher: Ecological Society of America.

[4] Billick, I. & Case, T. J. Higher Order Interactions in Ecological Communities: What Are They and How Can They be Detected? Ecology 75, 1530–1543 (1994). URL https://www.jstor.org/stable/1939614. Publisher: Ecological Society of America.

[5] Pomerantz, M. J. Do “Higher Order Interactions” in Competition Systems Really Exist? The American Naturalist 117, 583–591 (1981). URL https://www.jstor.org/stable/2460470. Publisher: [University of Chicago Press, American Society of Naturalists].

[6] Worthen, W. B. & Moore, J. L. Higher-Order Interactions and Indirect Effects: A Resolution Using Laboratory Drosophila Communities. The American Naturalist 138, 1092–1104 (1991). URL https://www.jstor.org/stable/2462509. Publisher: [University of Chicago Press, American Society of Naturalists].

[7] Morin, P. J., Lawler, S. P. & Johnson, E. A. Competition Between Aquatic Insects and Vertebrates: Interaction Strength and Higher Order Interactions. Ecology 69, 1401–1409 (1988). URL https://onlinelibrary.wiley.com/doi/abs/10.2307/1941637. _eprint: https://onlinelibrary.wiley.com/doi/pdf/10.2307/1941637.

[8] Battiston, F. et al. Networks beyond pairwise interactions: Structure and dynamics. Physics Reports (2020). URL http://www.sciencedirect.com/science/article/pii/S0370157320302489.

[9] Lambiotte, R., Rosvall, M. & Scholtes, I. From networks to optimal higher-order models of complex systems. Nature Physics 15, 313–320 (2019). URL https://www.nature.com/articles/s41567-019-0459-y. Number: 4 Publisher: Nature Publishing Group.

[10] Sanchez-Gorostiaga, A., Bajić, D., Osborne, M. L., Poyatos, J. F. & Sanchez, A. High-order interactions distort the functional landscape of microbial consortia. PLOS Biology 17, e3000550 (2019). URL https://journals.plos.org/plosbiology/article?id=10.1371/journal.pbio.3000550. Publisher: Public Library of Science.

[11] Tekin, E., Yeh, P. J. & Savage, V. M. General Form for Interaction Measures and Framework for Deriving Higher-Order Emergent Effects. Frontiers in Ecology and Evolution 6 (2018). URL https://www.frontiersin.org/articles/10.3389/fevo.2018.00166/full. Publisher: Frontiers.

[12] Yitbarek, S., Guittar, J. L., Knutie, S. A. & Ogbunugafor, C. B. Deconstructing higher-order interactions in the microbiota: A theoretical examination. preprint, Ecology (2019). URL http://biorxiv.org/lookup/doi/10.1101/647156.

[13] Mayfield, M. M. & Stouffer, D. B. Higher-order interactions capture unexplained complexity in diverse communities. Nature Ecology & Evolution 1, 1–7 (2017). URL https://www.nature.com/articles/s41559-016-0062. Number: 3 Publisher: Nature Publishing Group.

[14] Mickalide, H. & Kuehn, S. Higher-Order Interaction between Species Inhibits Bacterial Invasion of a Phototroph-Predator Microbial Community. Cell Systems 9, 521–533.e10 (2019). URL http://www.sciencedirect.com/science/article/pii/S2405471219303904.

[15] Sundarraman, D. et al. Higher-Order Interactions Dampen Pairwise Competition in the Zebrafish Gut Microbiome. mBio 11 (2020). URL https://mbio.asm.org/content/11/5/e01667-20. Publisher: American Society for Microbiology Section: Research Article.

[16] Chesson, P. Mechanisms of Maintenance of Species Diversity. Annual Review of Ecology and Systematics 31, 343–366 (2000). URL https://doi.org/10.1146/annurev.ecolsys.31.1.343. _eprint: https://doi.org/10.1146/annurev.ecolsys.31.1.343.

[17] Barabás, G., J. Michalska-Smith M. & Allesina, S. The Effect of Intra- and Interspecific Competition on Coexistence in Multispecies Communities. The American Naturalist 188, E1–E12 (2016). URL https://www.journals.uchicago.edu/doi/full/10.1086/686901. Publisher: The University of Chicago Press.

[18] Gardner, M. R. & Ashby, W. R. Connectance of Large Dynamic (Cybernetic) Systems: Critical Values for Stability. Nature 228, 784–784 (1970). URL https://www.nature.com/articles/228784a0. Bandiera_abtest: a Cg_type: Nature Research Journals Number: 5273 Primary_atype: Research Publisher: Nature Publishing Group.

[19] May, R. M. Will a Large Complex System be Stable? Nature 238, 413–414 (1972). URL https://www.nature.com/articles/238413a0. Bandiera_abtest: a Cg_type: Nature Research Journals Number: 5364 Primary_atype: Research Publisher: Nature Publishing Group.

[20] Allesina, S. & Tang, S. Stability criteria for complex ecosystems. Nature 483, 205–208 (2012). URL https://www.nature.com/articles/nature10832. Number: 7388 Publisher: Nature Publishing Group.

[21] Bunin, G. Ecological communities with Lotka-Volterra dynamics. Physical Review E 95, 042414 (2017). URL https://link.aps.org/doi/10.1103/PhysRevE.95.042414. Publisher: American Physical Society.

[22] Dougoud, M., Vinckenbosch, L., Rohr, R. P., Bersier, L.-F. & Mazza, C. The feasibility of equilibria in large ecosystems: A primary but neglected concept in the complexity-stability debate. PLOS Computational Biology 14, e1005988 (2018). URL https://journals.plos.org/ploscompbiol/article?id=10.1371/journal.pcbi.1005988. Publisher: Public Library of Science.

[23] Grilli, J. et al. Feasibility and coexistence of large ecological communities. Nature Communications 8, 14389 (2017). URL https://www.nature.com/articles/ncomms14389. Number: 1 Publisher: Nature Publishing Group.

[24] Stone, L. The feasibility and stability of large complex biological networks: a random matrix approach. Scientific Reports 8, 8246 (2018). URL https://www.nature.com/articles/s41598-018-26486-2/briefing/signup/. Bandiera_abtest: a Cc_license_type: cc_by Cg_type: Nature Research Journals Number: 1 Primary_atype: Research Publisher: Nature Publishing Group Subject_term: Ecological modelling;Ecological networks;Theoretical ecology Subject_term_id: ecological-modelling;ecological-networks;theoretical-ecology.

[25] Bairey, E., Kelsic, E. D. & Kishony, R. High-order species interactions shape ecosystem diversity. Nature Communications 7, 12285 (2016). URL https://www.nature.com/articles/ncomms12285. Number: 1 Publisher: Nature Publishing Group.

[26] Mézard, M., Parisi, G. & Virasoro, M. A. SK Model: The Replica Solution without Replicas. Europhysics Letters (EPL) 1, 77–82 (1986). URL https://doi.org/10.1209/0295-5075/1/2/006. Publisher: IOP Publishing.

[27] Diederich, S. & Opper, M. Replicators with random interactions: A solvable model. Physical Review A 39, 4333–4336 (1989). URL https://link.aps.org/doi/10.1103/PhysRevA.39.4333. Publisher: American Physical Society.

[28] Barbier, M. & Arnoldi, J.-F. The cavity method for community ecology. preprint, Ecology (2017). URL http://biorxiv.org/lookup/doi/10.1101/147728.

[29] Barbier, M., Arnoldi, J.-F., Bunin, G. & Loreau, M. Generic assembly patterns in complex ecological communities. Proceedings of the National Academy of Sciences 115, 2156–2161 (2018). URL https://www.pnas.org/content/115/9/2156. Publisher: National Academy of Sciences Section: Biological Sciences.

[30] Singh, P. & Baruah, G. Higher order interactions and species coexistence. Theoretical Ecology (2020). URL https://doi.org/10.1007/s12080-020-00481-8.

[31] AlAdwani, M. & Saavedra, S. Is the addition of higher-order interactions in ecological models increasing the understanding of ecological dynamics? Mathematical Biosciences 315, 108222 (2019). URL http://www.sciencedirect.com/science/article/pii/S0025556418307247.

[32] AlAdwani, M. & Saavedra, S. Ecological models: higher complexity in, higher feasibility out. Journal of The Royal Society Interface 17, 20200607 (2020). URL https://royalsocietypublishing.org/doi/10.1098/rsif.2020.0607. Publisher: Royal Society.

[33] Song, C., Rohr, R. P. & Saavedra, S. A guideline to study the feasibility domain of multi-trophic and changing ecological communities. Journal of Theoretical Biology 450, 30–36 (2018). URL https://www.sciencedirect.com/science/article/pii/S0022519318302042.

[34] Saavedra, S. et al. A structural approach for understanding multispecies coexistence. Ecological Monographs 87, 470–486 (2017). URL https://esajournals.onlinelibrary.wiley.com/doi/abs/10.1002/ecm.1263. _eprint: https://esajournals.onlinelibrary.wiley.com/doi/pdf/10.1002/ecm.1263.

[35] Sidhom, L. & Galla, T. Ecological communities from random generalized Lotka-Volterra dynamics with nonlinear feedback. Physical Review E 101, 032101 (2020). URL https://link.aps.org/doi/10.1103/PhysRevE.101.032101. Publisher: American Physical Society.

[36] Galla, T. Dynamically evolved community size and stability of random Lotka-Volterra ecosystems. EPL (Europhysics Letters) 123, 48004 (2018). URL https://doi.org/10.1209%2F0295-5075%2F123%2F48004. Publisher: IOP Publishing.

[37] Cui, W., Marsland III, R. & Mehta, P. Diverse communities behave like typical random ecosystems. 1904.02610 [cond-mat, physics:physics, q-bio] (2019). URL http://arxiv.org/abs/1904.02610. ArXiv: 1904.02610.

[38] Hu, J., Amor, D. R., Barbier, M., Bunin, G. & Gore, J. Emergent phases of ecological diversity and dynamics mapped in microcosms. Tech. Rep. (2021). URL https://www.biorxiv.org/content/10.1101/2021.10.28.466339v1. xCompany: Cold Spring Harbor Laboratory Distributor: Cold Spring Harbor Laboratory Label: Cold Spring Harbor Laboratory Section: New Results Type: article.

[39] Mehta, P. & Marsland, R. Cross-feeding shapes both competition and cooperation in microbial ecosystems. preprint, Ecology (2021). URL http://biorxiv.org/lookup/doi/10.1101/2021.10.10.463852.

[40] Bunin, G. Interaction patterns and diversity in assembled ecological communities. 1607.04734 [cond-mat, physics:physics, q-bio] (2016). URL http://arxiv.org/abs/1607.04734. ArXiv: 1607.04734.

[41] Barbier, M., de Mazancourt, C., Loreau, M. & Bunin, G. Fingerprints of High-Dimensional Coexistence in Complex Ecosystems. Physical Review X 11, 011009 (2021). URL https://link.aps.org/doi/10.1103/PhysRevX.11.011009.

[42] Jacquet, C. et al. No complexity–stability relationship in empirical ecosystems. Nature Communications 7, 12573 (2016). URL https://www.nature.com/articles/ncomms12573. Number: 1 Publisher: Nature Publishing Group.

[43] Gause, G. Experimental Studies on the Struggle for Existence: I. Mixed Population of Two Species of Yeast (1932). URL /paper/Experimental-Studies-on-the-Struggle-for-Existence%3A-Gause/06cc528cc8951a76afa74e6ed65e52f9603cd9ea.

[44] Letten, A. D. & Stouffer, D. B. The mechanistic basis for higher-order interactions and non-additivity in competitive communities. Ecology Letters 22, 423–436 (2019). URL https://onlinelibrary.wiley.com/doi/abs/10.1111/ele.13211. _eprint: https://onlinelibrary.wiley.com/doi/pdf/10.1111/ele.13211.

[45] Momeni, B., Xie, L. & Shou, W. Lotka-Volterra pairwise modeling fails to capture diverse pairwise microbial interactions. eLife 6, e25051 (2017). URL https://doi.org/10.7554/eLife.25051. Publisher: eLife Sciences Publications, Ltd.

[46] Soetaert, K., Petzoldt, T. & Setzer, R. W. Solving Differential Equations in R: Package deSolve. Journal of Statistical Software 33, 1–25 (2010). URL https://doi.org/10.18637/jss.v033.i09.

[47] Wilson, W. G. et al. Biodiversity and species interactions: extending Lotka–Volterra community theory. Ecology Letters 6, 944–952 (2003). URL https://onlinelibrary.wiley.com/doi/abs/10.1046/j.1461-0248.2003.00521.x. _eprint: https://onlinelibrary.wiley.com/doi/pdf/10.1046/j.1461-0248.2003.00521.x.

[48] Wilson, W. G. & Lundberg, P. Biodiversity and the Lotka–Volterra theory of species interactions: open systems and the distribution of logarithmic densities. Proceedings of the Royal Society of London. Series B: Biological Sciences 271, 1977–1984 (2004). URL https://royalsocietypublishing.org/doi/10.1098/rspb.2004.2809. Publisher: Royal Society.

