## Supplementary Information for "Coexistence in diverse communities with higher-order interactions"

### 1 Model Form

We consider the same model as in Eq. 1 of the main text. The dynamics of species $i$ are given by

$$\dot{N}_i = N_i \left( R_i - d_i N_i - \sum_{j \neq i} A_{ij} N_j - \sum_j N_j \sum_k B_{ijk} N_k \right) \tag{S1}$$

and all parameters have the same interpretation as in the main text. Note that, in this model, the effect of species $j$ on the interaction between $k$ and $i$, $B_{ikj}$, also multiplies the product of the abundances of species $j$ and $k$, but is a separate interaction in our model. We sample species growth rates and interaction parameters from probability distributions with specified summary statistics. Our theory applies to arbitrary probability distributions (with mild assumptions on their moments) because it appeals to the central limit theorem, but we use normal distributions in our simulations. Specifically, we let

| *Growth Rates* | *Pairwise Interactions* | *Higher-Order Interactions* |
|---|---|---|
| $\langle R_i \rangle = \mu_R$ | $\langle A_{ij} \rangle = \mu_A / S$ | $\langle B_{ijk} \rangle = \mu_B / S^2$ |
| $\langle R_i^2 \rangle - \langle R_i \rangle^2 = \sigma_R^2$ | $\langle A_{ij}^2 \rangle - \langle A_{ij} \rangle^2 = \sigma_A^2 / S$ | $\langle B_{ijk}^2 \rangle - \langle B_{ijk} \rangle^2 = \sigma_B^2 / S^2$ |
| | $\langle A_{ij} A_{ji} \rangle - \langle A_{ij} \rangle^2 = \rho_A \sigma_A^2 / S$ | |

which is the same as in the main text, except now each off-diagonal pairwise interaction coefficient $A_{ij}$ is correlated to its partner reflected across the diagonal $A_{ji}$ through the parameter $\rho_A$. We concentrate on systems which are competitive on average, so that $\mu_A, \mu_B \geq 0$ but the coefficients themselves, $A_{ij}$ and $B_{ijk}$, can be positive or negative since they are sampled from distributions. Because there are many more higher-order interactions than pairwise interactions, the summed effect of the former will often exceed that of the latter. This combinatorial effect is not of particular interest, so we scaled the interaction statistics to account for the number of possible interactions. As a result, the mean pairwise interaction strength that a focal species experiences is $\mu_A$, because it experiences pairwise interactions from $S$ species, while the mean higher-order interaction strength that a focal species experiences is $\mu_B$. We investigate the coexistence properties of the model without these scalings in the last set of results in the main text.

### 2 Cavity Method

In this section, we provide the cavity calculation in detail and then derive some of its consequences. Our calculation is a fairly standard application of the methodology [1] and it closely follows the calculations in recent related work [2–4]. Until the Finite size corrections section, we neglect all finite size corrections in the calculation.

#### 2.1 Cavity calculation

We first re-write Equation S1 using the statistics of the random interactions:

$$\begin{aligned}\frac{dN_i}{dt} &= N_i\left(R_i - N_i - \sum_{j\neq i} A_{ij}N_j - \sum_j N_j \sum_k B_{ijk}N_k\right)\\ &= N_i\left(\mu_R + \sigma_R r_i - N_i - \frac{\mu_A}{S}\sum_{j\neq i} N_j + \frac{\sigma_A}{\sqrt{S}}\sum_{j\neq i} a_{ij}N_j - \frac{\mu_B}{S^2}\sum_j\sum_k N_jN_k + \frac{\sigma_B}{S}\sum_j N_j \sum_k b_{ijk}N_k\right)\\ &= N_i\left(K_i - N_i + \frac{\sigma_A}{\sqrt{S}}\sum_{j\neq i} a_{ij}N_j + \frac{\sigma_B}{S}\sum_j N_j\sum_k b_{ijk}N_k\right)\end{aligned} \tag{S2}$$

where we have defined $K_i = \mu_R + \sigma_r r_i - \frac{\mu_A}{S}\sum_{j\neq i} N_j - \frac{\mu_B}{S^2}\sum_j\sum_k N_jN_k$ and a new set of random variables $r_i$, $a_{ij}$ and $b_{ijk}$ with the statistics $\langle r_i\rangle = \langle a_{ij}\rangle = \langle b_{ijk}\rangle = 0$, $\langle r_i^2\rangle = \langle a_{ij}^2\rangle = \langle b_{ijk}^2\rangle = 1$ and $\langle a_{ij}a_{ji}\rangle = \rho_A$. In deriving Equation S2, we have dropped the $O(1/S)$ corrections that arise from not allowing pure self-regulation in both the pairwise and higher-order interaction terms. There is still self-regulation both through the $N_i^2$ term and the higher-order interactions with coefficients $b_{iij}$ and $b_{iji}$ for all $i, j$. Throughout the calculation, we neglect corrections that vanish in the $S \to \infty$ limit (but see the Finite size corrections section). By re-writing the model into Equation S2, we have isolated the effect of different interaction statistics on the ecological dynamics.

Suppose the system has reached an equilibrium where some species coexist. If species $i$ has positive abundance at this equilibrium, then its abundance is given by

$$N_i = K_i + \frac{\sigma_A}{\sqrt{S}}\sum_{j\neq i} a_{ij}N_j + \frac{\sigma_B}{S}\sum_j N_j\sum_k b_{ijk}N_k\ . \tag{S3}$$

We now add an invading species which we will index species $0$ and approximate species $i$'s abundances after the invasion. Specifically, we add a perturbation

$$\delta_i = N_0\left(\frac{\sigma_A}{\sqrt{S}}a_{i0} + \frac{\sigma_B}{S}\sum_j N_{j\backslash 0}\left(b_{i0j} + b_{ij0}\right) + \frac{\sigma_B}{S}b_{i00}N_0\right) \tag{S4}$$

which is the new set of interactions that species $0$ introduces into species $i$'s dynamics. Here, $N_{j\backslash 0}$ denotes the equilibrium abundance of species $j$ before the introduction of species $0$. Adding $\delta_i N_i$

to each $\dot{N}_i$ and Taylor-expanding the response of each species to its perturbation, we find that:

$$\begin{aligned} N_i &= N_{i\setminus 0} + \sum_j v_{ij}\delta_j \\ &= N_{i\setminus 0} + N_0 \sum_j v_{ij} \left( \frac{\sigma_A}{\sqrt{S}} a_{j0} + \frac{\sigma_B}{S} \sum_k N_{k\setminus 0} \left(b_{j0k} + b_{jk0}\right) + \frac{\sigma_B}{S} b_{j00} N_0 \right) \\ &= N_{i\setminus 0} + N_0 L_i \end{aligned} \tag{S5}$$

where $v_{ij} = \left.\frac{\partial N_i}{\partial \delta_j}\right|_{\delta_j = 0}$ is the response of species $i$ to perturbation $j$ and we have defined

$$L_i = \sum_j v_{ij} \left( \frac{\sigma_A}{\sqrt{S}} a_{j0} + \frac{\sigma_B}{S} \sum_k N_{k\setminus 0} \left(b_{j0k} + b_{jk0}\right) + \frac{\sigma_B}{S} b_{j00} N_0 \right) . \tag{S6}$$

We have derived these expressions assuming that the abundance of the invader is at some fixed $N_0$, but we do not know the value of $N_0$. In fact, it will in turn depend on the expressions for $N_i$. More concretely, we have expressed the abundances after a perturbation in terms of the abundances before the perturbations and the abundances of the invading species. We substitute these expressions for the abundances of the non-invading species into the formula for species $0$ and re-arrange to solve for species $0$'s abundances:

$$\begin{aligned} N_0 &= K_0 + \frac{\sigma_A}{\sqrt{S}} \sum_{j\neq 0} a_{0j} \left(N_{j\setminus 0} + N_0 L_j\right) + \frac{\sigma_B}{S} \sum_j \left(N_{j\setminus 0} + N_0 L_j\right) \sum_k b_{0jk} \left(N_{k\setminus 0} + N_0 L_k\right) \\ &= \left[ K_0 + \frac{\sigma_A}{\sqrt{S}} \sum_{j\neq 0} a_{0j} N_{j\setminus 0} + \frac{\sigma_B}{S} \sum_j \sum_k b_{0jk} N_{j\setminus 0} N_{k\setminus 0} \right] \\ &\quad + N_0 \left[ \frac{\sigma_A}{\sqrt{S}} \sum_{j\neq 0} a_{0j} L_j + \frac{\sigma_B}{S} \sum_j \sum_k b_{0jk} \left(N_{j\setminus 0} L_k + N_{k\setminus 0} L_j\right) \right] \\ &\quad + N_0^2 \left[ \frac{\sigma_B}{S} \sum_j \sum_k b_{0jk} L_j L_k \right] . \end{aligned} \tag{S7}$$

In Equation S7, we have treated the abundances of the non-invading community members as fixed (once they have responded to the invasion). As a result, Equation S7 approximately solves for the abundance that the invader can achieve while accounting for its own influence on the community composition. It is not exact,

In contrast to the pairwise case, we have found a quadratic equation for species $0$'s abundance. However, the quadratic coefficient of this equation is mean zero since $\langle b_{0jk} \rangle = 0$ and $b_{0jk}$ is independent of all the terms in any $L_i$. Therefore, we get that

$$\left\langle \frac{\sigma_B}{S} \sum_j \sum_k b_{0jk} L_j L_k \right\rangle = \frac{\sigma_B}{S} \sum_j \sum_k \langle b_{0jk} \rangle \langle L_j L_k \rangle = 0$$

and we have, on average, a linear equation for species $0$'s abundance. Following [2–4], we can now solve for species $0$'s abundance

$$N_0 = \frac{K_0 + \frac{\sigma_A}{\sqrt{S}} \sum_{j \neq 0} a_{0j} N_{j\backslash 0} + \frac{\sigma_B}{S} \sum_j \sum_k b_{0jk} N_{j\backslash 0} N_{k\backslash 0}}{1 - \frac{\sigma_A}{\sqrt{S}} \sum_{j \neq 0} a_{0j} L_j + \frac{\sigma_B}{S} \sum_j \sum_k b_{0jk} \left( N_{j\backslash 0} L_k + N_{k\backslash 0} L_j \right)} \tag{S8}$$

and approximate the denominator. First, we note that

$$\begin{aligned} \langle a_{0i} L_i \rangle &= \left\langle a_{0i} \sum_j v_{ij} \left( \frac{\sigma_A}{\sqrt{S}} a_{j0} + \frac{\sigma_B}{S} \sum_k N_{k\backslash 0} \left( b_{j0k} + b_{jk0} \right) + \frac{\sigma_B}{S} b_{j00} N_0 \right) \right\rangle \\ &= \left\langle \frac{\sigma_A}{\sqrt{S}} \sum_j v_{ij} a_{0i} a_{j0} \right\rangle = \left\langle \frac{\sigma_A}{\sqrt{S}} v_{ii} a_{0i} a_{i0} \right\rangle \end{aligned} \tag{S9}$$

since $a_{0i}$ is independent of all the HOI terms and every pairwise term except for $a_{i0}$. Using Equation S9, we find that

$$\frac{\sigma_A}{\sqrt{S}} \sum_{j \neq 0} a_{0j} L_j \approx \left\langle \frac{\sigma_A^2}{S} \sum_{j \neq 0} v_{jj} a_{0j} a_{j0} \right\rangle + O\left( \frac{1}{\sqrt{S}} \right) = v \rho_A \sigma_A^2 \phi + O\left( \frac{1}{\sqrt{S}} \right) \tag{S10}$$

where we have defined $v = \sum_i v_{ii}$ to be the average response of a species to its own perturbation and $\phi$ to be the fraction of species that coexist in the equilibrium that species $0$ is invading. Using arguments similar to the those that let us reduce Equation S7 to a linear equation, we find that the HOI terms in the denominator of Equation S8 only contribute fluctuations which go to zero in the limit $S \to \infty$, so we neglect them. All in all, we have a new formula for species $0$'s abundance:

$$N_0 = \frac{K_0 + \frac{\sigma_A}{\sqrt{S}} \sum_{j \neq 0} a_{0j} N_{j\backslash 0} + \frac{\sigma_B}{S} \sum_j \sum_k b_{0jk} N_{j\backslash 0} N_{k\backslash 0}}{1 - v \rho_A \sigma_A^2 \phi} . \tag{S11}$$

Now, we analyze the numerator of Equation S11. It is the sum of many weakly correlated random variables, so we expect it to be well-approximated by a Gaussian from the central limit theorem. The mean and variance of this Gaussian are simple functions of the fraction of coexisting species $\phi$, the first two moments of the coexisting species ($\langle N \rangle$ and $\langle N^2 \rangle$) and the underlying interaction statistics. For the mean of the Gaussian, we get that

$$\begin{aligned}\left\langle K_0 + \frac{\sigma_A}{\sqrt{S}}\sum_{j\neq 0} a_{0j}N_{j\backslash 0} + \frac{\sigma_B}{S}\sum_j\sum_k b_{0jk}N_{j\backslash 0}N_{k\backslash 0}\right\rangle \\ = \mu_R + \sigma_r\langle r_i\rangle - \frac{\mu_A}{S}\sum_{j\neq i} N_j - \frac{\mu_B}{S^2}\sum_j\sum_k N_jN_k \\ = \mu_R - \mu_A\phi\langle N\rangle - \mu_B\phi^2\langle N\rangle^2\end{aligned} \tag{S12}$$

where we have neglected the $O\left(\frac{1}{S}\right)$ corrections from the missing self-regulation terms. Using the fact that species growth rates, pairwise interactions and higher-order interaction are all independent, we compute the variance of the Gaussian as

$$\begin{aligned}\text{Var}\left[K_0 + \frac{\sigma_A}{\sqrt{S}}\sum_{j\neq 0} a_{0j}N_{j\backslash 0} + \frac{\sigma_B}{S}\sum_j\sum_k b_{0jk}N_{j\backslash 0}N_{k\backslash 0}\right] \\ = \text{Var}\left[K_0\right] + \frac{\sigma_A^2}{S}\left\langle\left(\sum_{j\neq 0} a_{0j}N_{j\backslash 0}\right)^2\right\rangle + \frac{\sigma_B^2}{S^2}\left\langle\left(\sum_j\sum_k N_{j\backslash 0}N_{k\backslash 0}b_{0jk}\right)^2\right\rangle \\ = \sigma_R^2 + \sigma_A^2\phi\langle N^2\rangle + \sigma_B^2\phi^2\langle N^2\rangle^2\ .\end{aligned} \tag{S13}$$

There are corrections to the variance that come from the higher-order interactions which provide self-regulation to other species mediated through species $0$. These corrections will depend on the fourth moment of the species abundance distribution $\langle N^4\rangle$. However, these corrections are again order $O(1/S)$ so we neglect them.

All together, we have now determined the distribution of invader abundances. Specifically, we have found that the mean and variance of the invader abundances are

$$\mu_0 = \frac{\mu_R - \mu_A\phi\langle N\rangle - \mu_B\phi^2\langle N\rangle^2}{1 - v\rho_A\sigma_A^2\phi} \qquad \text{and} \qquad \sigma_0^2 = \frac{\sigma_R^2 + \sigma_A^2\phi\langle N^2\rangle + \sigma_B^2\phi^2\langle N^2\rangle^2}{(1 - v\rho_A\sigma_A^2\phi)^2} \tag{S14}$$

respectively. Then, the invader abundances follow a Guassian distribution with mean $\mu_0$ and variance $\sigma_0^2$ which we will denote by $N_0 \sim P(N_0|\,\mu_0, \sigma_0^2)$. If we sample from this distribution and get a positive abundance, then the invader will achieve this abundance and establish in the community. If instead we get a negative abundance, then species $0$ will be unable to invade and its abundance will go to zero. The denominator of our formulas for the mean and variance of this distribution account for the feedback that the invasion of species $0$ has on its own growth. If the pairwise interactions are uncorrelated (ie. $\rho_A = 0$), then $N_0$ is determined only by the community state before invasion. In this case, we could have derived our formula for $N_0$ by guessing that each $N_i$ at equilibrium is an independent realization of the same Gaussian distribution and then computing its mean and variance directly from the feasibility condition in Equation S3, as we stated in the main text. In fact, earlier studies did this calculation for the Lotka-Volterra model without appealing to the cavity method [5, 6].

Last, we conclude that the distribution of successful invaders is the same as the distribution of coexisting species. Therefore, if we restrict the distribution of invader abundances to be positive values, we have actually found the species abundance distribution (SAD) for our randomly parametrized model. By relating the distribution of $N_0$ to the distribution of coexisting species, we also derive coupled equations for the fraction of coexisting species $\phi$, the mean community abundance $\langle N \rangle$ and the second moment of the SAD $\langle N^2 \rangle$:

$$\begin{aligned} \phi &= \int_0^\infty P(N_0 | \, \mu_0, \sigma_0^2) \; dN_0 \\ \langle N \rangle &= \frac{1}{\phi} \int_0^\infty N_0 P(N_0 | \, \mu_0, \sigma_0^2) \; dN_0 \\ \langle N^2 \rangle &= \frac{1}{\phi} \int_0^\infty N_0^2 P(N_0 | \, \mu_0, \sigma_0^2) \; dN_0 \end{aligned} \tag{S15}$$

where the factors of $1/\phi$ normalizes the integral. The last unknown parameter in the problem is $v$, and we can derive a fourth equation using the definition of $v_{ij}$ and by differentiating Equation S8 with respect to the perturbation $\delta_i$. We get that

$$v = \frac{1}{1 - v \rho_A \sigma_A^2 \phi} \tag{S16}$$

and we now have four equations in four unknowns that we can solve numerically to determine the species abundance distribution. Our derivation is non-rigorous – it is based on a self-consistent calculation of the statistics of the species abundance distribution. Moreover, we have made several approximations throughout the calculation. We must check our predictions against numerical simulations, as we do in the main text and in the Simulation Results section.

#### 2.2 Solving the system of equations

We solve the system of nonlinear equations in Eq. S15 and Eq. S16 numerically for the specified growth rate and interaction statistics to get predictions for $\phi$, $\langle N \rangle$ and $\langle N^2 \rangle$. In solving these equations, we change variables so that we actually integrate over the standard normal distribution.

As a result, we can re-write Eq. S15 using the error function $\mathrm{erf}$ as

$$\begin{aligned}
\phi &= \frac{1}{2}\left(1+\mathrm{erf}\left(\frac{\mu_0}{\sqrt{2}\sigma_0}\right)\right)\\
\langle N\rangle &= \frac{1}{\phi}\left[\frac{\mu_0}{2}\left(1+\mathrm{erf}\left(\frac{\mu_0}{\sqrt{2}\sigma_0}\right)\right)+\frac{\sigma_0}{\sqrt{2\pi}}\exp\left(-\frac{\mu_0^2}{2\sigma_0^2}\right)\right]\\
\langle N^2\rangle &= \frac{1}{\phi}\left[\frac{\mu_0^2+\sigma_0^2}{2}\left(1+\mathrm{erf}\left(\frac{\mu_0}{\sqrt{2}\sigma_0}\right)\right)+\frac{\mu_0\sigma_0}{\sqrt{2\pi}}\exp\left(-\frac{\mu_0^2}{2\sigma_0^2}\right)\right]\ .
\end{aligned} \tag{S17}$$

From these equations, it is clear that the ratio $\mu_0/\sigma_0$ plays a crucial role in the predictions for $\phi$, $\langle N\rangle$ and $\langle N^2\rangle$. Specifically, when $\sigma_0$ is small so that $\mu_0/\sigma_0$ is large, $\mathrm{erf}\left(\frac{\mu_0}{\sqrt{2}\sigma_0}\right)=1$ because it integrates over the entire normal distribution. In this case, $\phi \approx 1$ and all species coexist in a community with a finite number of species. A similar logic applies to the effect of $\mu_0$ on coexistence, as we describe in the main text.

#### 2.3 The $\phi = 1$ limit when $\rho_A = 0$

It is difficult to understand the system of equations in Eq. S17 intuitively. In this section, we set $\rho_A = 0$ and consider the limit where all species coexist so that we can build intuition. As we described in the main text, the cavity equations simplify considerably in this limit because $\mu_0 = \langle N\rangle$ and $\sigma_0^2 = \langle N^2\rangle - \langle N\rangle^2$. Once we have solved for $\mu_0$ and $\sigma_0$, we can then compute the ratio $\mu_0/\sigma_0$ and determine how it behaves as a function of the growth rate or interaction statistics. In general, if $\mu_0/\sigma_0$ increases (resp. decreases) as a function of one of the growth rate or interaction statistics, then this statistic increases (resp. decreases) the fraction of coexisting species. In the pairwise case (ie. when $\mu_B = \sigma_B = 0$), we find that the ratio is

$$\left.\frac{\mu_0}{\sigma_0}\right|_{\mu_B=\sigma_B=0} = \sqrt{\frac{\mu_R^2\left(1-\sigma_A^2\right)}{\sigma_A^2(1-\mu_A)^2+\mu_R^2\sigma_A^2}}\ . \tag{S18}$$

When $\sigma_R = 0$, we find that this ratio only depends on $\sigma_A$, and it decreases as $\sigma_A$ increases. Even when $\sigma_R \neq 0$, $\sigma_A$ has the same qualitative effect on coexistence. Otherwise, increasing $\mu_R$ increase this ratio, while increasing $\mu_A$ decreases it, as we saw in Fig. 3 of the main text. Now, we compute this ratio when there are higher-order interactions in the system, though we now leave it in terms of $\mu_0$ so it is easier to analyze. We find that

$$\frac{\mu_0}{\sigma_0} = \sqrt{\frac{2\mu_0^2\sigma_B^2}{1-\sigma_A^2-2\mu_0^2\sigma_B^2-\sqrt{(1-\sigma_A^2)^2-4\sigma_B^2(\sigma_R^2+\mu_0^2)}}} \tag{S19}$$

which is not particularly illuminating, so we plot its behavior as a function of $\mu_0$ for a few different choices of $\sigma_R$ in Fig. A. We find the exact same dependence as in the full cavity method equations. First, let's note that $\mu_0$ increases as $\mu_R$ increase, and decreases as $\mu_A$ or $\mu_B$ increase. When $\sigma_R = 0$, the ratio decreases when $\mu_0$ increases, meaning that increasing the mean abundance of the species actually decreases the fraction of coexisting species. When $\sigma_R$ is large, the ratio increases as $\mu_0$ increases, so that more species coexist when the average abundance is larger. There are intermediate values of $\sigma_R$ where the ratio first increases and then decreases, as our results in the main text suggest.

#### 2.4 Effect of mean interaction strength when $\sigma_R = 0$

In the main text and in the cavity calculation, we showed theoretically that the mean pairwise interaction strength has no affect on coexistence when all species had the same growth rate ($\sigma_R = 0$) and the number of species is large ($S \to \infty$). In fact, the mean pairwise interaction strength can have a small effect on coexistence in our simulations due to our parameterization of the model in Eq. S1. Because we set $A_{ii} = 0$ in S1, changing the mean pairwise interaction strength does not change the level of self-regulation in the community. As a result, there is a small bias in our simulation results, where communities with more competitive interactions exhibit lower fractions of coexisting species (Fig. D). In Fig. B, we show that this bias disappears as $S$ increases. In other words, it is an effect of the finite size of the simulations. Moreover, we can show analytically that, if $A_{ii} = \mu_A/S$, then this bias no longer exists. The basic intuition is that, when $A_{ii} = \mu_A/S$, changing the mean interaction strength changes both all of the competitive interactions and the degree of self-regulation in exactly the same way. As a result, the mean abundance may change, but no species can be excluded, because the structure of the interactions has not changed. Specifically, when there are no higher-order interactions, the equilibrium where all species coexist solves $\vec{R} = [dI + A]\vec{N}$ for some matrix $A$. Let's assume that $N_i > 0$ for all $i$ and let's guess that the new equilibrium abundance vector is given by $\kappa\vec{N}$ for some constant $\kappa$ when we replace $A$ by $A + \mu_A/S\vec{1}\vec{1}^T$ where $\vec{1}$ denotes the vector of all 1's. Then, we have that

$$\begin{aligned}\vec{R} &= [dI + A + \mu_A/S\vec{1}\vec{1}^T]\kappa\vec{N} \\ &= \kappa\vec{R} + \mu_A\kappa\langle N\rangle\vec{1}\end{aligned} \tag{S20}$$

and, since $\vec{R} = r\vec{R}$ for some positive constant growth rate $r$, Eq. S20 simply becomes $r = \kappa r + \mu_A\kappa\langle N\rangle \implies \kappa = \frac{r}{r+\mu_A\langle N\rangle}$. This $\kappa$ solution is positive whenever the interactions are on average competitive $\mu_A > 0$ and becomes negative when the interactions become too mutualistic to converge to equilibrium. In Fig. B, we provide numerical evidence of this argument, by adjusting the diagonal entries $d_i$ to be $\mu_A$. In fact, essentially the same argument shows that the mean interaction strength has no effect on coexistence for higher-order interactions when there is mean-adjusted cubic self-regulation (see the Cubic self-regulation and pure competition section).

##### 2.5 Finite size corrections

Throughout the cavity method calculation, we dropped terms that disappeared in the $S \to \infty$ limit. In our simulations, the number of species is necessarily finite, and so it is possible that these neglected terms could prove important in shaping the fraction of species that coexist, especially in systems with relatively small numbers of species. To this end, we now state modified versions of Eq. S14 where we have included corrections for the finite system size. We have that

$$\mu_0 = \frac{\mu_R - \mu_A \phi \langle N \rangle - \left(1 - \frac{1}{S}\right) \mu_B \phi^2 \langle N \rangle^2}{(1 - \mu_A/S)(1 - v\rho_A \sigma_A^2 \phi)}$$
$$\sigma_0^2 = \frac{\sigma_R^2 + \left(1 - \frac{1}{S}\right) \sigma_A^2 \phi \langle N^2 \rangle + \left(1 - \frac{1}{S}\right) \sigma_B^2 \phi^2 \langle N^2 \rangle^2}{(1 - \mu_A/S)^2 (1 - v\rho_A \sigma_A^2 \phi)^2} \ . \qquad \text{(S21)}$$

The main effect of these alterations is through the term in the denominator which corrects for the fact that $A_{ii} = 0$ in our parameterization. In particular, these terms incorporate the effect of mean interaction strength discussed in the previous section into the cavity method framework, and they account for the small differences in the fraction of coexisting species in Fig. 3(A) of the main text. In our simulations, we also observe small differences due to the effect of mean pairwise interaction strength (see Fig. B and D). Throughout this work, we specify a number of species $S$ even when we are only plotting cavity method predictions and solve the system of equations specified by Eq. S21 numerically. See the code at https://github.com/tgibbs-hub/CavityHOIs for the details of the numerical solving procedure.

#### 3 Simulation Results

In this section, we describe the results of additional numerical simulations. The code to run our simulations and generate our figures is available on GitHub at
https://github.com/tgibbs-hub/CavityHOIs.

##### 3.1 Additional simulations to test cavity method predictions

In Fig. C, we plot the results of simulations and the predictions from the cavity method for the fraction of coexisting species, the average species abundance and the variance in species abundances when there are $S = 300$ species in the system. We consider three different possible parameterizations of the interactions. In the pairwise case, there are only pairwise interactions, while in the higher-order interaction case, there are only higher-order interactions. In the mixed case, we set both mean pairwise and higher-order interaction strengths as $\mu_A = \mu_B = 2$. We also choose to increment the variations in pairwise and higher-order interaction variabilities equally so that the total variability $\sqrt{\sigma_A^2 + \sigma_B^2}$ matches that of the pairwise or higher-order interaction variability. In these simulations, the predictions work quite well. In fact, there are significantly fewer replicates in these cases as compared to the simulations with smaller numbers of species, but the predictions seem more accurate. The data is most variable in the higher-order case, but this is because a relatively large fraction of species coexist here. When there are only higher-order interactions, the pairwise correlations have no effect since there are no pairwise interactions.

In Fig. D and E, we plot the same predictions as in Fig. 3 of the main text, but now we include simulation data for $100$ replicates of $S = 30$ species in all cases. We find that the predictions work quite well even at this relatively small number of species. In Fig. D, we also plot the predicted and simulated mean and variance in abundances of the coexisting species.

In Fig. F, we solve the cavity method equations for a large range of different mean interaction strengths and variabilities in interaction strengths. We consider systems with purely pairwise and purely higher-order interactions, as well as mixtures of the two as described in the previous paragraph. In the pairwise and mixed cases, the correlation in pairwise interactions ($\rho_A$) has a negative affect on coexistence, with anti-correlated interactions promoting coexistence [2, 3]. We set all species growth rates to be the same value ($\sigma_R = 0$), so the mean pairwise interaction strength $\mu_A$ has an imperceptible effect on the fraction of coexisting species when there are only pairwise interactions because there are $S = 300$ species. However, we can see from the differing areas that the red region covers in these plots that the mean interaction strength does affect the ability of our algorithm to find a numerical solution. In fact, $\mu_A$ can affect the dynamics of the Lotka-Volterra model (even when $\sigma_R = 0$) by controlling whether or not the strength of competition is large enough to prevent mutualisms from giving rise to diverging species abundances (as is signified by the red area). When there are higher-order interactions, the species abundances can diverge more easily, as shown by the larger red areas in Fig. F. This is likely because these mutualistic interactions are now proportional to the square of the interacting species, meaning that they have an even greater potential for fast mutual growth. In the mixed and purely higher-order cases, we also see the effect of mean competition strength that we documented in the main text – fewer species coexist as the mean competition strength becomes smaller.

#### 3.2 Multiple attracting equilibria and divergence in abundances

The cavity method is exact when there is an uninvasible, unique and attracting fixed point of the dynamics, but its predictions can still be accurate when there are multiple attractors [2, 3]. Despite the fact that in both the pairwise and higher-order interaction parametrizations it is possible for there to be multiple stable fixed points, we did not observe them in the simulations we ran. So, even though the higher-order interaction system can have many more fixed points (even up to $S$ feasible fixed points [7]) than a pairwise community, these extra fixed points are likely not having a large impact on the dynamics in the parameter regimes we have explored. Specifically, we solve the dynamics of a given set of interaction strengths multiple times and compute the average difference between species abundances between these replicates. We also average over different realizations of the interaction strengths for the same interaction statistics ($\sigma_A$ or $\sigma_B$ in Fig. G). In all of these simulations, we did not observe a non-zero average difference. We did observe, however, simulations in which the dynamics did not reach equlibrium. In these simulation runs, the species abundances grew indefinitely. We only observed the divergence of species abundances when there are purely higher-order interactions, but it is possible for pairwise interactions, as shown in [2, 3] and predicted in Fig. F. In Fig. G, as $\sigma_B$ increased, the probability of reaching equilibrium decreased. Throughout this work, we exclude cases from our simulation results where the abundances diverged. The divergence in species abundances is in part caused by having a smaller numbers of species in the system. For more diverse systems, the value of $\sigma_A$ or $\sigma_B$ at which the divergence in species abundances first occurs becomes more sharply defined [2].

#### 3.3 Feasibility controls when the first species is excluded

We now simulate the dynamics of of interaction networks with the same structure, but different statistics, to understand how the species abundances themselves change as a function of the interaction statistics. To accomplish this, we specify an interaction network (pairwise or higher-order) by sampling the required interactions from a standard normal distribution. Then, we multiply all these coefficients by the desired interaction variation ($\sigma_A$ or $\sigma_B$) and add the desired competition strength ($\mu_A$ or $\mu_B$). In this way, we can create a variety of different networks with the same structure, but different statistics (as in [8, 9]). We then numerically solve the dynamics of each of these networks starting at different initial conditions to observe how individual species abundances change as a function of the interaction statistics (Fig. H). In fact, this is the same procedure we use to find the critical interaction variations from the main text, except here we are still scaling the interaction statistics as in our standard parameterizations. Specifically, the variation in interaction strengths at which the first species becomes excluded can be read off of the x-axis in each of the panels of Fig. H.

If there were multiple attracting equilibria, the curves in Fig. H would not be continuous functions of the interaction statistics. Similarly, it is clear from Fig. H that the dynamics are attracted to the "same" fixed point as we change the variation in interaction strengths. This fixed point does not lose stability as the variation in interaction strengths increases. Instead, some species decrease in abundance as the variation in interaction strengths increases, and eventually are excluded. Interestingly, this result suggests that we need only consider the feasibility of the fixed point where all species coexist, and not its stability, to determine the critical interaction variability we defined in the main text. More generally, it seems that predicting the behavior of this fixed point as a function of the variation in interaction strengths, even when all species do not coexist, is enough to predict the simulated abundances.

##### 3.4 High abundance species and mutualisms

In our cavity method calculation, we showed that the ration $\mu_0/\sigma_0$ plays a key role in determining the fraction of species that coexist. We then derived the same qualitative dependence of the fraction of coexisting species on the mean competition strength that we observed in simulations and the full cavity method from the limit where pairwise interactions are uncorrelated and all species coexist. We now explicitly construct the ratio $\mu_0/\sigma_0$ from our simulated abundance data and from the full cavity method solution. The data and predictions match quite well (Fig. I). When this ratio is larger, fewer species coexist, and we find that the simulated and predicted ratios are consistent with the coexistence results presented in the main text. Using the reasoning described in Methods section of the main text, the cavity method formalism explains the counter-intuitive effect of mean competition strength on coexistence, but it does not provide an intuitive understanding, at the level of species interactions, of how weaker competition produces fewer coexisting species.

The interactions that remain in the community after the it has assembled are not a random subset of the interactions initially present. Instead, they are biased to promote coexistence [10, 11], because species that experience more competitive interaction strengths are more likely to have smaller abundances and eventually be excluded. In fact, there is a fairly direct relationship between the interaction strengths that a species experience (ie. $\sum_j A_{ij}$ or $\sum_j \sum_k B_{ijk}$ for species $i$) and that species' abundance, especially when all species coexist (Fig. J). When some species are excluded, this correspondence is not as precise but it still exists.

These patterns, however, do not fully describe the impact of interactions on species growth, because they do not incorporate the species abundance involved in the interactions. To address this, we now ask whether or not more (resp. less) abundant species interact in a characteristic pattern with the other more (resp. less) abundant species in the community. In Fig. K(A) (resp. Fig. K(B)), we plot the interaction strengths ($A_{ij}$ for pairwise interactions and $B_{ijk}$ for higher-order interactions) of the most (resp. least) abundant species in the community as a function of the abundances that engage in that interaction ($N_j$ for pairwise interactions and $N_jN_k$ for higher-order interactions). We can quantify the tendency of a given species to experience mutualistic interactions from high abundance species by the slope of a regression on these data. If this slope is positive, then species tend to experience competitive interactions from more abundant species, because the value of the interaction strength increases as a function of the species abundances involved in that interaction. Conversely, if the slope is negative, then species tend to experience more mutualistic interactions from the most abundant species, while experiencing more competitive interactions from the least abundant species. These regression slopes tend to become more negative as a function of the focal species' abundance, suggesting that more abundant species interact mutualistically with one another, but more competitively with lower abundance species (see Fig. K(C)). This behavior has been documented previously [10, 11], and it is likely to be a general phenomenon for community assembly in ecological models. In our model, where higher-order interactions depend on the square of species abundances, this bias in interaction strengths creates abundant cliques of species which suppress other species in the community (see Fig. H). When in addition $\mu_B$ is small, these abundant cliques form strong mutualisms between their members, further increasing their abundances and driving many species to exclusion.

If species interactions saturate, rather than grow indefinitely, as a function of species abundances, then mutualisms between the dominant species may not produce the same effect of mean competition strength on coexistence. In the main text, we showed that introducing saturation eliminated the counter-intuitive effect of mean competition strength. Here, we provide plots of the ratio between the mean and standard deviation in coexisting species (Fig. L) which display the same qualitative behavior as the coexistence plots in the main text, suggesting that saturation successfully dampened the growth of the variance as a function of species mean abundance. In this case, we are plotting the ratio mean to standard deviation of the truncated species abundance distribution ie. the species abundance distribution without negative abundances. In previous plots (Fig. A and I), we plotted this ratio for the non-truncated species abundance distribution, because this is the quantity that actually enters the cavity method calculation. Therefore, the behavior in Fig. L does not have the same theoretical justification as in Fig. A and I, but it is nonetheless reassuring that it is consistent with the coexistence results.

#### 3.5 Cubic self-regulation and pure competition

In the main text and the previous section, we argued that the strong mutualistic interactions created cliques of highly abundant species that exclude many other species, specifically when $\mu_B$ is small and $\sigma_R$ or $\sigma_B$ is large. In a model with saturating interactions, the strength of these mutualistic interactions can no longer grow indefinitely, and we showed in simulations that increasing the mean competition strength always decreased the fraction of coexisting species in this case. In the dynamics of two competing species, however, if the strength of the mutualistic interactions exceeds the strength of self-regulation, then the species abundances diverge. In modeling saturating interactions, we limited the strength of the inter-specific mutualisms, thereby promoting coexistence. However, we also could have left the inter-specific interactions unmodified and instead increased the strength of intra-specific competition to counteract the strong mutualistic higher-order interactions. To this end, we now simulate a model where species experience cubic, rather than quadratic, self-regulation. Specifically, the dynamics of species $i$ is now given by

$$\dot{N}_i = N_i \left( R_i - d_i N_i^2 - \sum_j N_j \sum_k B_{ijk} N_k \right) \tag{S22}$$

so that there are no pairwise interactions. We consider two different parameterizations of the $d_i$ parameter analogous to those we analyzed in the Effect of mean interaction strength when $\sigma_R = 0$ section. We either set $d_i = 1$ or $d_i = 1 + \mu_B / S^2$ so that changing the mean higher-order interaction strength affects the all higher-order terms, including the self-regulatory ones.

In both cases, when there is no variability in growth rates ($\sigma_R = 0$) the effect of mean higher-order interaction strength on coexistence is very small throughout a range of variations in interaction strengths (see Fig. M). When species have different growth rates, systems with more competitive interactions have fewer coexisting species (see Fig. M). This is consistent with both the results for purely pairwise systems, as well as systems with pairwise and saturating higher-order interactions. In the case where we adjust the cubic self-regulation parameter using the mean higher-order interaction strength, the same argument that we presented in the Effect of mean interaction strength when $\sigma_R = 0$, except with $\kappa = \sqrt{\frac{r}{r + \mu_B \langle N^2 \rangle}}$, shows that the mean higher-order interaction strength does not affect which species coexist. In other words, this case is precisely consistent with the effect of the mean interaction strength in the pairwise case.

We identified strong mutualistic interactions as the source of the counter-intuitive effect of mean higher-order interaction strength on coexistence. A natural test of this hypothesis is to simply eliminate mutualistic interactions from the community (for example, by truncating the distribution of interaction coefficients at zero) and then observe whether or not more competitive higher-order interactions still produce a higher fraction of coexisting species. There is a complication with this approach, however, because constraining the distribution of interaction coefficients to be positive introduces a relationship between the mean and variance of the distribution. Therefore, these parameters cannot be independently varied as we have done throughout this work. In addition, we need to consider our choice of scalings for the pairwise and higher-order interaction statistics to ensure that mutualistic interactions are not allowed. In Fig. N, we plot the predicted fraction of coexisting species from the cavity method where we have constrained the statistics of the interactions to be the same as those from a uniform distribution between $0$ and some maximum interaction strength. In these predictions, we are simultaneously varying the mean and variance of the interactions so that there are no mutualistic interactions. Increasing the maximum interaction strength (and hence both the mean interaction strength and the variance in interaction strengths) decreases the fraction of species that coexist (Fig. N). Overall, this behavior is consistent with our interpretation of our results – less competitive higher-order interactions reduce coexistence relative to more competitive ones because strong mutualistic interactions cause there to be dominant cliques of species which suppress the others. When we prevent mutualistic interactions, this behavior also disappears.

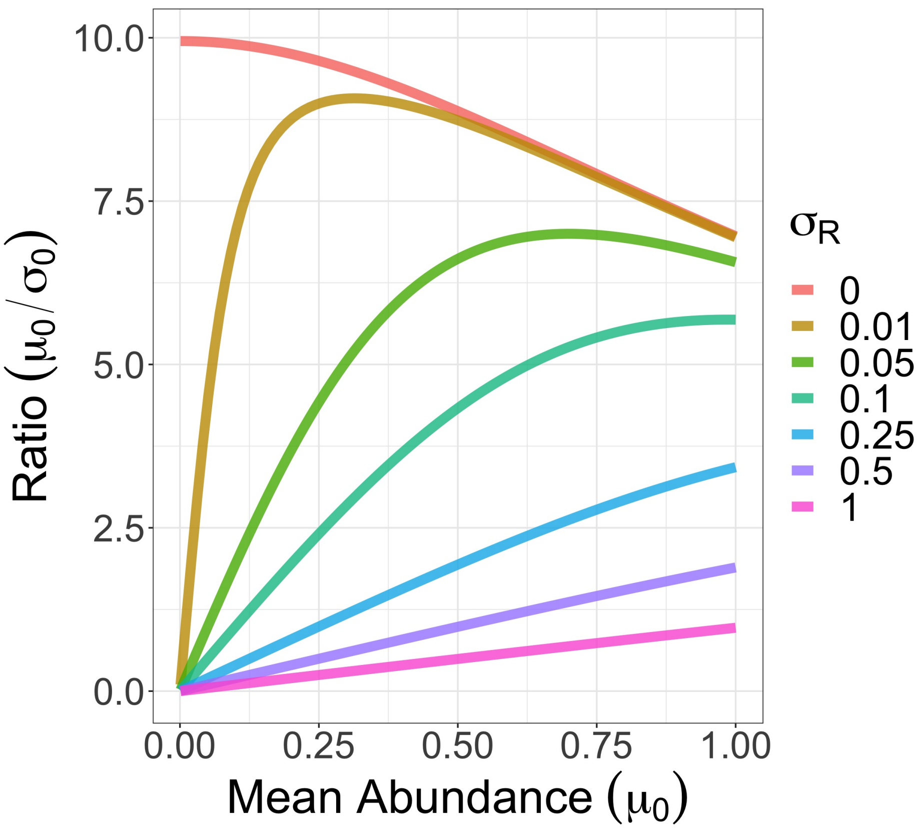

Figure A: **The ratio $\mu_0/\sigma_0$ as a function of $\mu_0$.** We plot the predicted ratio $\mu_0/\sigma_0$ from Eq. S19 as we vary $\mu_0$ for a few different choices of $\sigma_R$. When $\sigma_R = 0$, this ratio always decreases as a function of $\mu_0$, showing that communities with smaller average abundance actually have more species coexisting. When $\sigma_R \neq 0$, there is always a regime in which the opposite dependence is true. For large enough $\mu_0$ though, we recover the same dependence as when $\sigma_R = 0$.

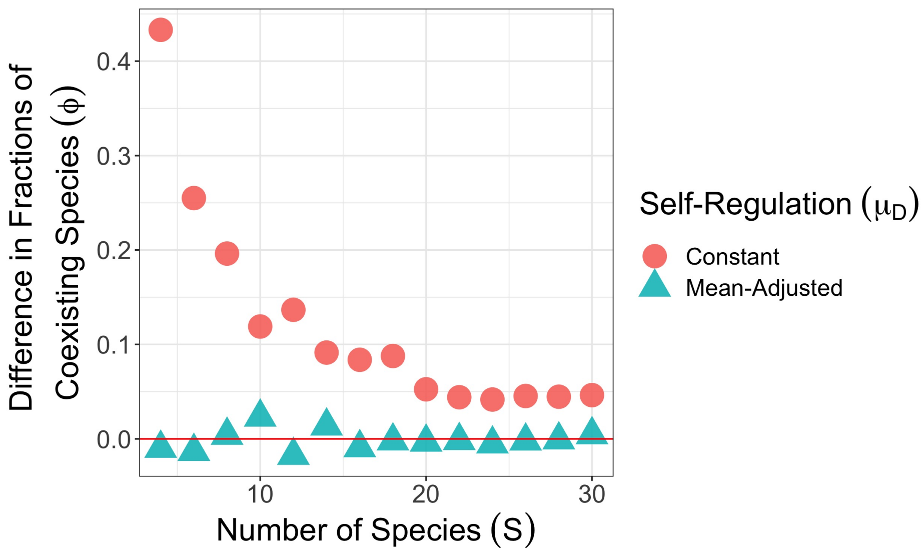

Figure B: **Effect of pairwise mean interaction strength when species have equal growth rates.** In each point, we compute the difference between the fraction of coexisting species when $\mu_A = 1$ and $\mu_A = 4$ as we vary the number of species ($S$). A positive difference indicates that more species coexisted when $\mu_A = 1$, while a negative difference indicates that more species coexisted when $\mu_A = 4$. We average the fractions of coexisting species over 100 replicates. In the constant self-regulation case (red circles), we set $d_i = 1$, as we did throughout the main text, while in the mean-adjusted self-regulation case (blue triangles), we set $d_i = 1 - \mu_A/S$, so that the strength of self-regulation changes in the same way as the mean pairwise interaction strength. When we adjust the self-regulation by the mean competition strength, the difference in the fractions of coexisting species fluctuates around zero, indicating that there is no effect of mean pairwise interaction strength. When we do not adjust the self-regulation, increasing mean competition strength decreases coexistence, but this effect disappears as the number of species becomes large. For all points, we set $\mu_R = 1.5$, $\sigma_R = 0$ and $\sigma_A = 0.6$.

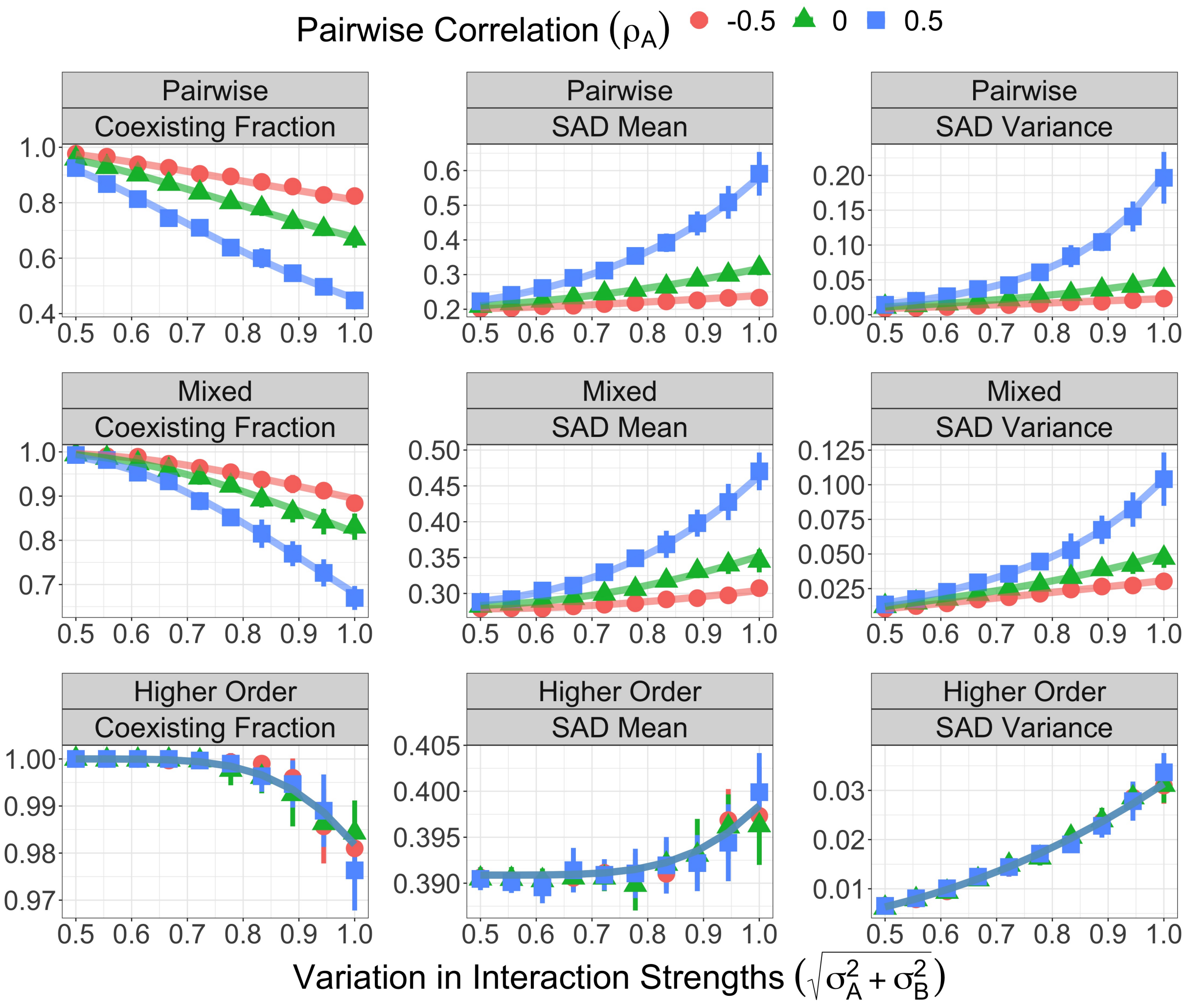

Figure C: **Cavity method predictions and simulation results for more diverse communities.** We plot the predictions from the cavity method (lines), the average of simulation results (shapes) and error bars showing one standard deviation as a function of the overall standard deviation of pairwise and higher-order interactions (ie. $\sqrt{\sigma_A^2+\sigma_B^2}$). The different colors shapes designate the different levels of pairwise correlation $\rho_A = -0.5, 0, 0.5$. There are 10 replicates for each of the correlated cases and 20 replicates for the for uncorrelated cases. Each panel is labeled by both the type of interactions in the community (purely pairwise interactions, a mixture of pairwise and higher-order interactions, and purely higher-order interactions) and the interaction statistic that is shown in the graph (coexisting fraction $\phi$, mean of the species abundance $\langle N \rangle$ and variance in the species abundances $\langle N^2 \rangle - \langle N \rangle^2$). In all cases, the number of species is $S = 300$, the total competition strength is $\mu_A + \mu_B = 4$ and all growth rates are $R_i = 1$ (ie. $\sigma_R = 0$ and $\mu_R = 1$).

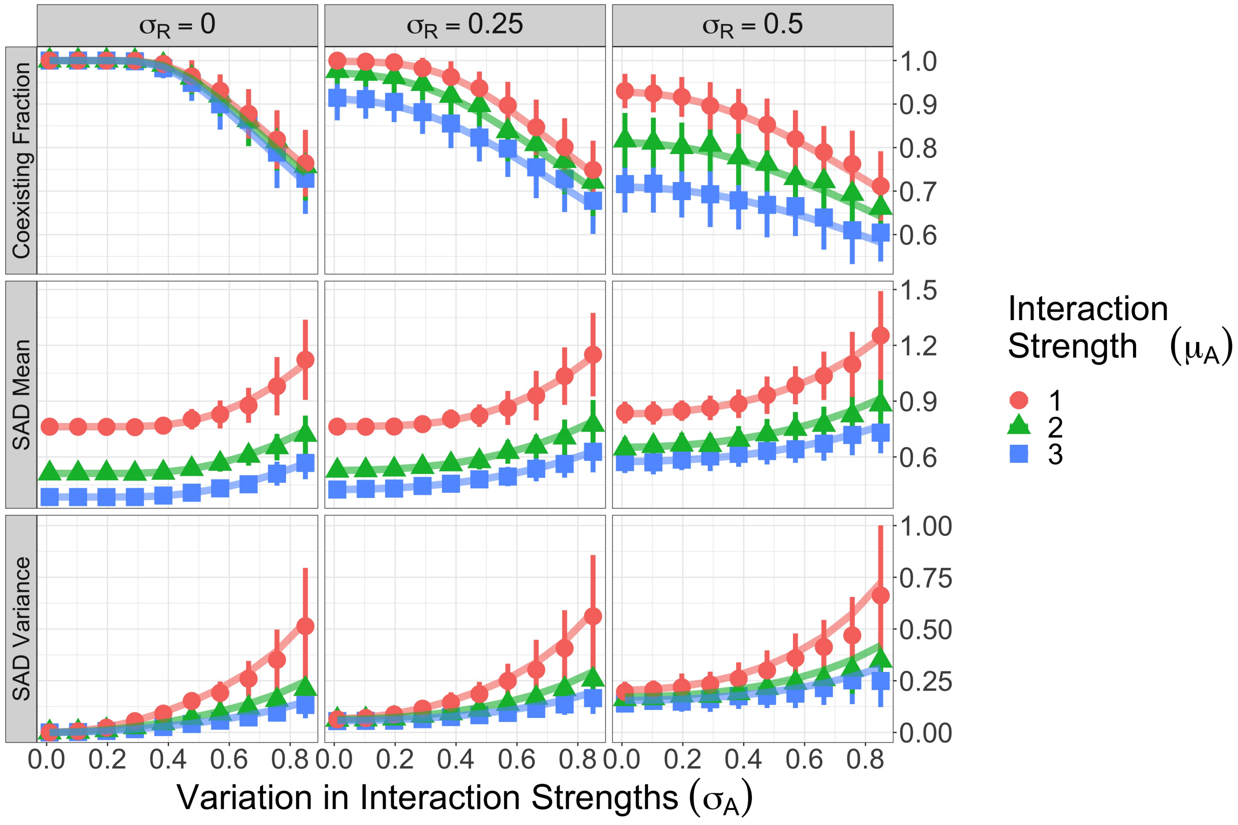

Figure D: **Simulation results for the pairwise case in Fig. 3 of the main text.** We plot the predictions from the cavity method (lines), the average of simulation results (shapes) and error bars showing one standard deviation as a function of the variability in pairwise interactions ($\sigma_A$). The rows are different macroecological properties ((coexisting fraction $\phi$, mean of the species abundance $\langle N \rangle$ and variance in the species abundances $\langle N^2 \rangle - \langle N \rangle^2$) while the columns are different variabilities in growth rates ($\sigma_R = 0, 0.25, 0.5$). The colors and shapes designate different levels of mean pairwise interaction strength ($\mu_A$). In all panels, $S = 30$ and $\mu_R = 1.5$. There are $S = 30$ species and each point represents 100 replicates.

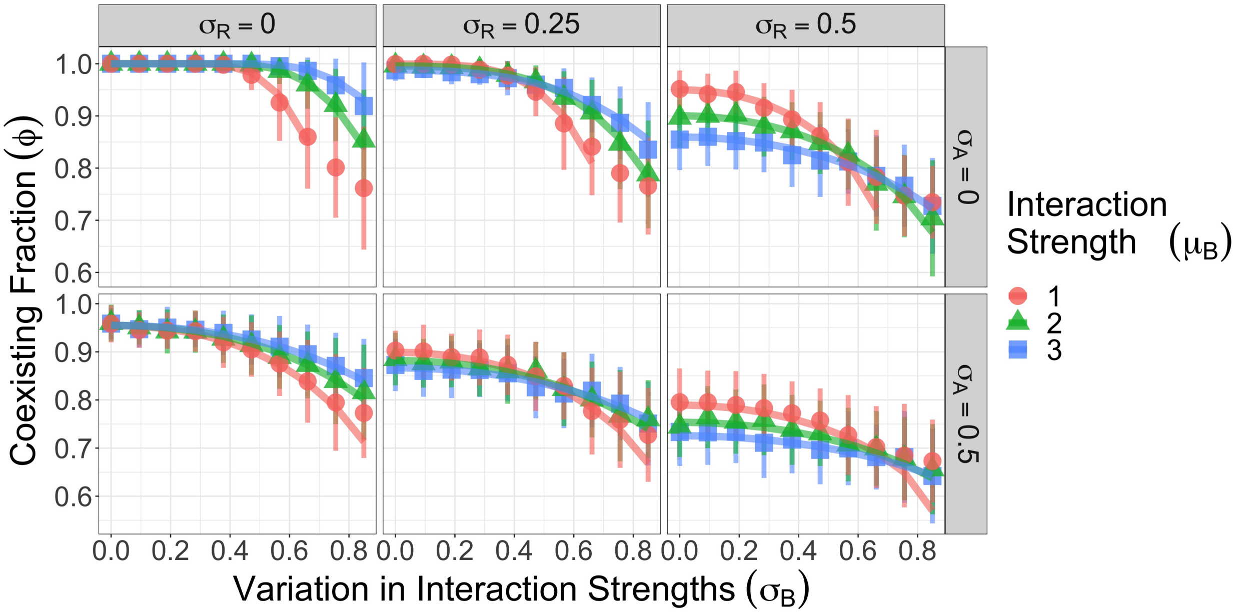

Figure E: **Simulation results for the higher-order cases in Fig. 3 of the main text.** We plot the predictions from the cavity method (lines), the average of simulation results (shapes) and error bars showing one standard deviation for the fraction of coexisting species ($\phi$) as a function of the variability in higher-order interactions ($\sigma_B$). The rows are different variabilities in the pairwise interactions ($\sigma_A = 0$ when there are no pairwise interactions and $\sigma_A = 0.5$ when there are pairwise interactions) while the columns are different variabilities in growth rates ($\sigma_R = 0, 0.25, 0.5$). In the second row (when there are pairwise interactions), the mean pairwise interaction strength is $\mu_A = 1$. The colors and shapes designate different levels of mean higher-order interaction strength ($\mu_B$). In all panels, $S = 30$ and $\mu_R = 1.5$. There are $S = 30$ species and each point represents 100 replicates.

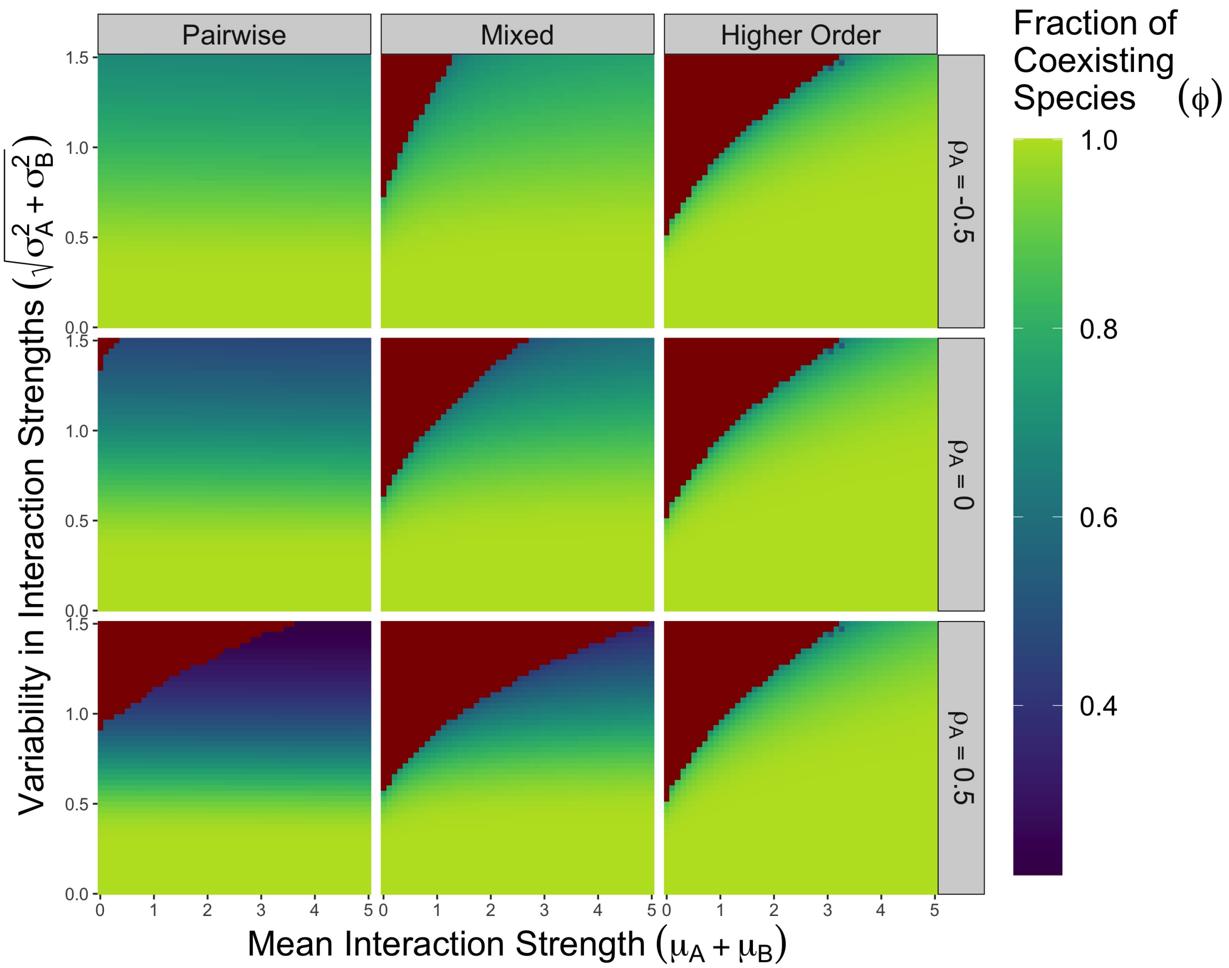

Figure F: **Cavity method predictions for the fraction of coexisting species** We plot the predicted fraction of coexisting species as a function of the overall mean interaction strength $(\mu_A + \mu_B)$ and the overall variability in interaction strengths $(\sqrt{\sigma_A^2 + \sigma_B^2})$. The different columns designate different types of interactions (purely pairwise interactions, a mixture of pairwise and higher-order interactions, and purely higher-order interactions), while the rows show different levels of pairwise correlation $(\rho_A = -0.5, 0, 0.5)$. Red colors indicate that our algorithm did not find a numerical solution, suggesting that the dynamics do not converge (see Fig. G and [2, 3]). We derive these predictions for a community of $S = 300$ species, effectively eliminating the effect of the number of species.

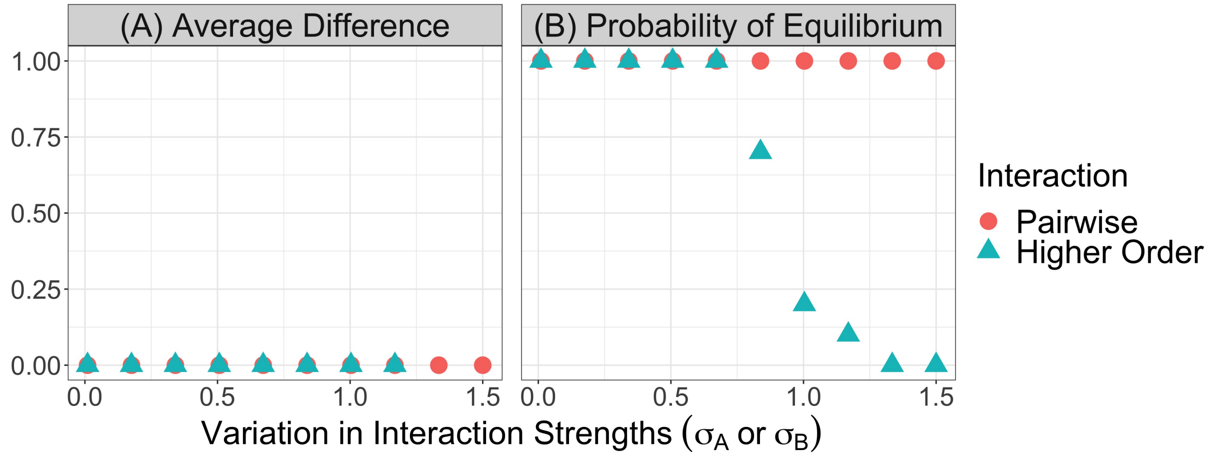

Figure G: **Diverging abundances and multiple equilibria** (A) We plot the average difference between the equilibria reached from different initial conditions for both pairwise and higher-order interactions. We run simulations of the same parameters with different initial conditions and different realizations of the parameters with the same statistics. We average over each species' abundance, the different initial conditions and the different replicates of the parameters, but we do not find any examples of multiple stable equilibria. If the dynamics do not reach equilibrium, we remove the points (as in the two higher-order interaction points with the largest variations in interaction strengths). (B) We plot the probability of finding an equilibrium (as defined in the main text) while we change the variation in interaction strengths for both pairwise and higher-order interactions. We compute the probability over the different initial conditions and the different replicates of parameters. When the higher-order interaction dynamics do not reach an equilibrium, it is because the abundances diverge. In both panels, we run 10 replicates of each point with 5 different randomly sampled initial conditions. We set $S = 50$, $\mu_R = 1.5$ and $\sigma_R = 0.25$ in all cases. For the pairwise interactions, we set $\mu_A = 2$, and for the higher-order interactions, we set $\mu_B = 2$.

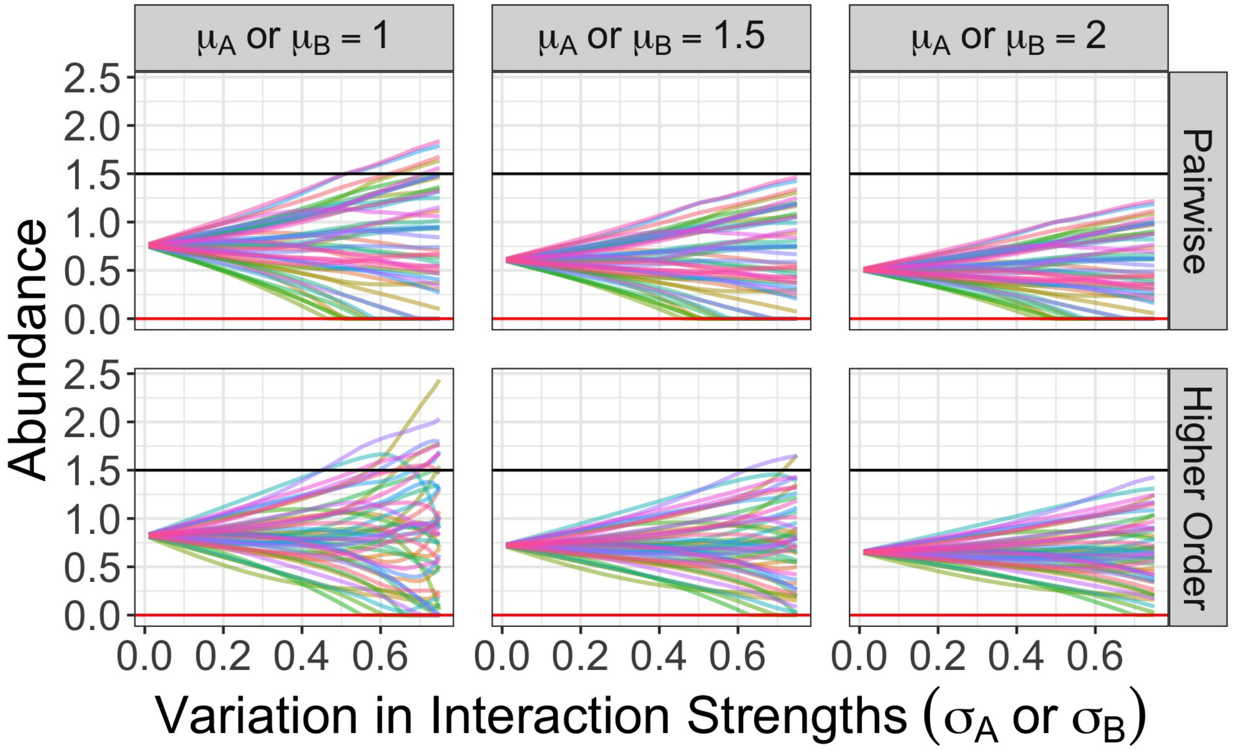

Figure H: **Species abundances as a function of interaction variation** We plot the species abundances while we vary the variation in interaction strengths for a single realization of the random pairwise and higher-order networks. Specifically, we sample the pairwise and higher-order interactions from a standard normal distribution to create the initial networks, and then we multiply by the specified interaction variability and add a constant mean interaction strength to form a network with the correct statistics. Each point on the curves above is a the equilibrium of a simulation with different initial conditions and different interaction statistics but the same underlying network structure. As a result, we can observe how individual species abundances change as the interaction variation increases for a few different possible mean competition strengths. In all panels, $S = 50$, $\mu_R = 1.5$ and $\sigma_R = 0$. The black lines in each panel denotes the species' carrying capacity in the absence of interactions, while the red line denotes zero abundance. Therefore, species with abundances exceeding the black line are experiencing a net mutualistic effect of all of their interactions.

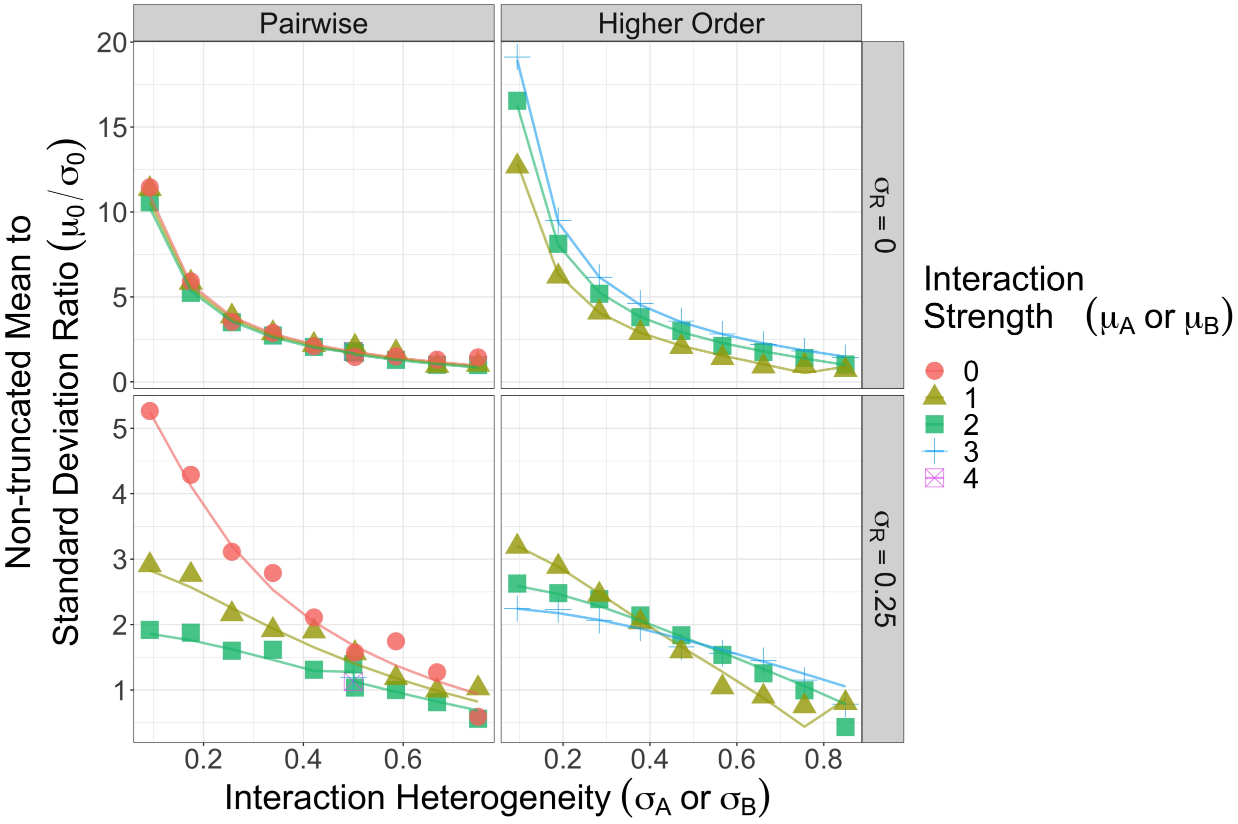

Figure I: **Simulated and predicted non-truncated mean-standard deviation ratios** We plot the ratio $\mu_0/\sigma_0$ (which we discussed when carrying out the cavity method calculation) using our simulations (shapes) and predictions (lines) for purely pairwise and purely higher-order interactions and two different variations in species growth rates. Larger values of this ratio indicate that fewer species will coexist. Parameters are the same as in Fig 3 of the main text or Fig. D and E.

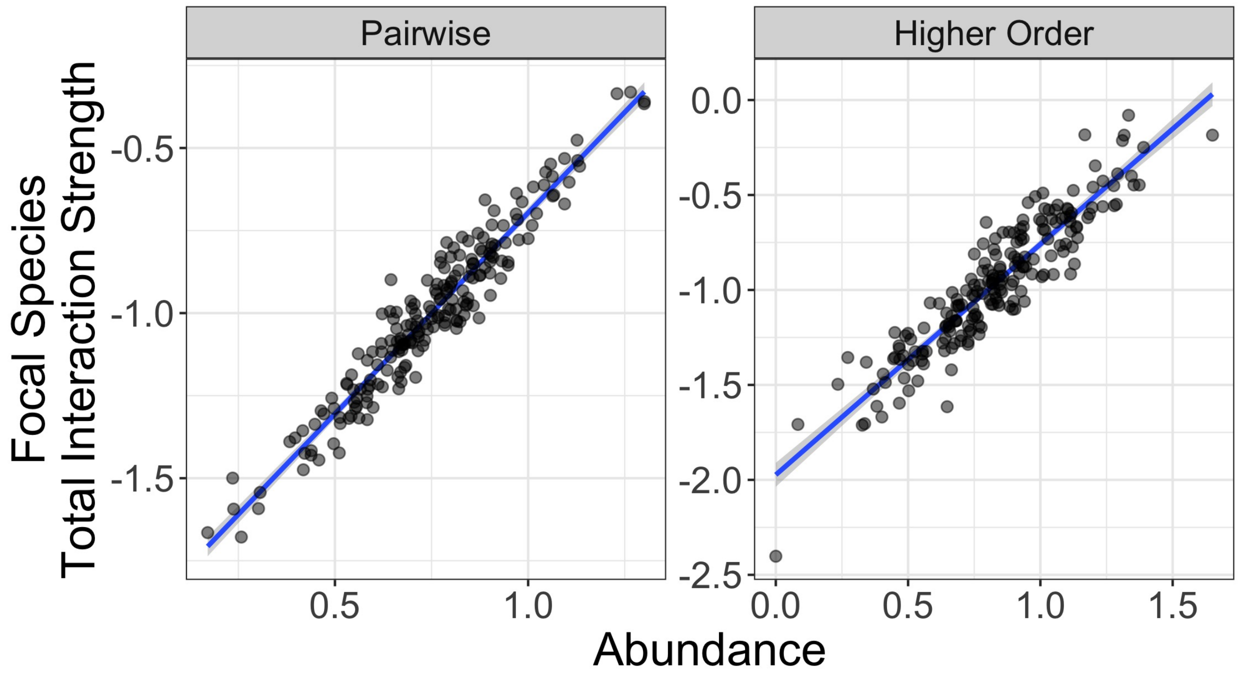

Figure J: **Relationship between total interaction strengths and abundance** For purely pairwise and purely higher-order interactions, we plot the total interaction strength a species experiences (ie. $\sum_j A_{ij}$ or $\sum_j \sum_k B_{ijk}$ for species $i$) as a function of the focal species' abundance. Blue lines are regressions with gray confidence intervals. The parameters are $S = 200$, $\mu_R = 1.5$, $\sigma_R = 0$, $\mu_A = \mu_B = 1$, $\sigma_A = 0.25$ and $\sigma_B = 0.35$.

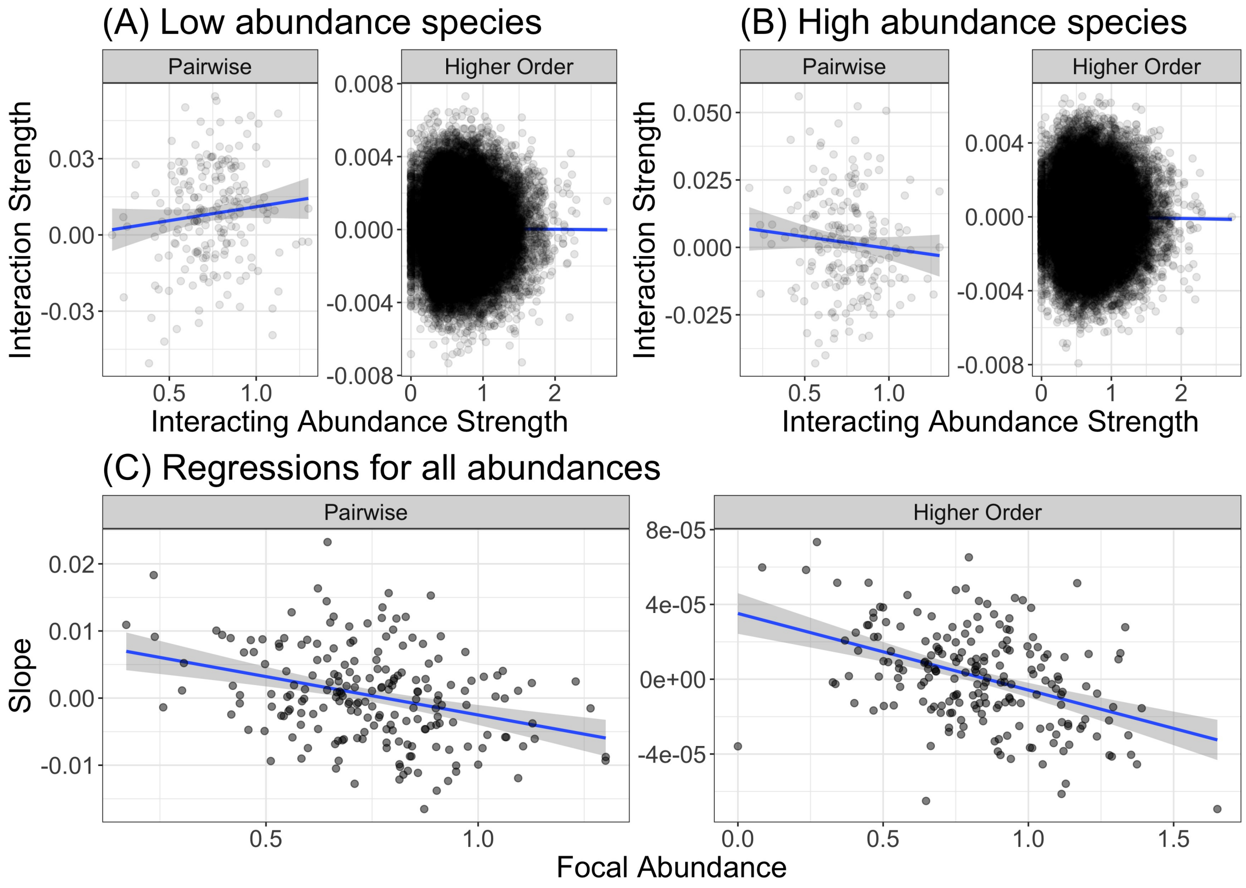

Figure K: **Species with large abundances engage in mutualisms with one another** (A) We plot, for both pairwise and higher-order interactions, the value of the interaction strength as a function of the abundances that create that interaction (ie. if $A_{ij}$ (resp. $B_{ijk}$) is the value on the y-axis, then $N_j$ (resp. $N_j N_k$) is the value on the x-axis) for the species $i$ with the smallest abundance in the community at equilibrium. (B) The same plot as in panel A but now for the species $i$ with the largest abundance at equilibrium. (C) We plot the slope of the regression lines (like those in panels A and B) for all species as a function of that species abundance in the coexisting community. We see that, for both pairwise and higher-order interactions, dominant species are more likely to interact mutualistically with other dominant species, while competing more strongly with species that have smaller abundances. The opposite pattern is true for species that are competitively suppressed at equilibrium. In all panels, blue lines are regressions with gray confidence intervals. The parameters are the same as in Fig. J.

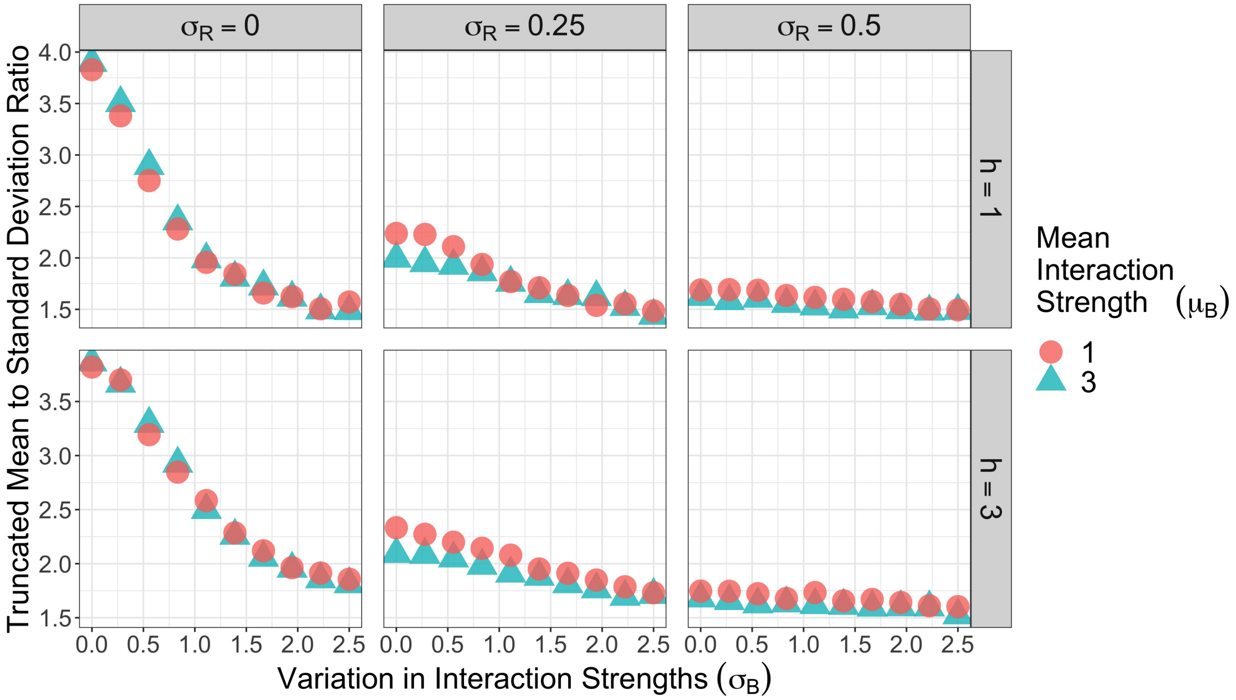

Figure L: **Mean to standard deviation ratio in simulated abundances for saturating higher-order interactions** We plot the ratio of the mean of the observed abundances ($\langle N \rangle$) to the standard deviation of the observed abundances ($\sqrt{\langle N^2 \rangle - \langle N \rangle^2}$) as a function of the variation in higher-order interaction strengths for a few different mean interaction strengths. The panels designate different variations in growth rates and levels of saturation. Importantly, this is not exactly the same ratio as $\mu_0/\sigma_0$ but it does show the same qualitative dependence that we observed in Fig. 4 of the main text – when interaction saturate strongly, this ratio is larger for the smaller value of the mean competition strength, suggesting that fewer species coexist at equilibrium. Parameters are the same as in Fig. 4 of the main text.

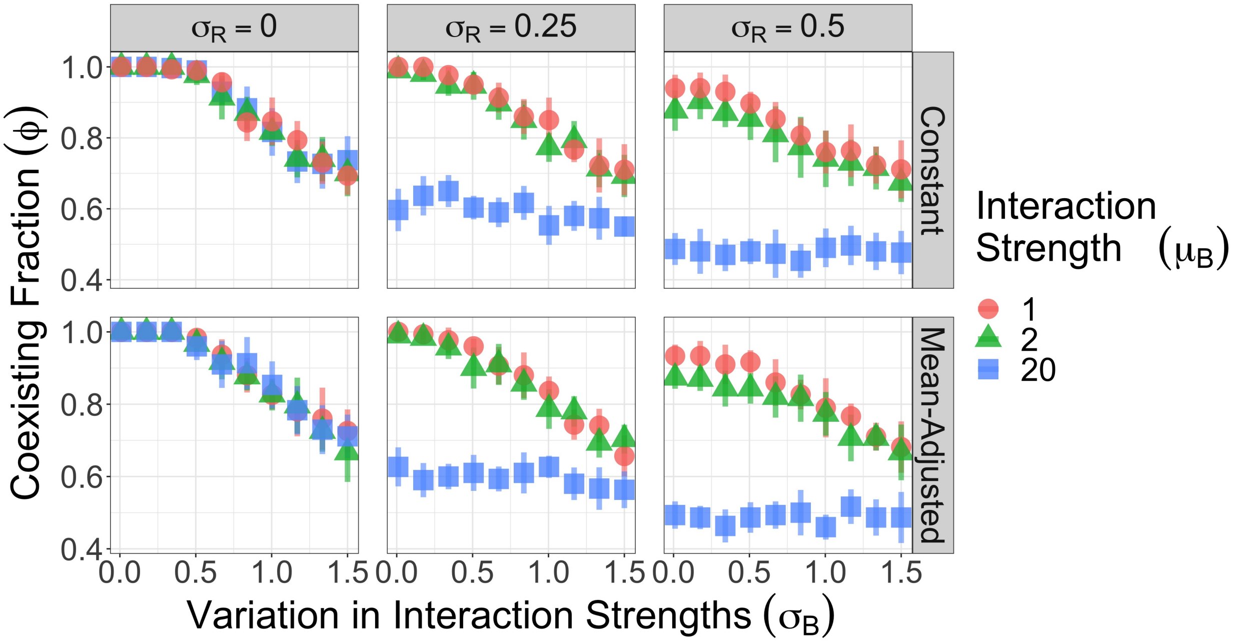

Figure M: **Coexistence with cubic self-regulation** We plot the fraction of species that coexist as a function of the variation in interaction strengths ($\sigma_B$) for the model in Eq. S22 where self-regulation is cubic rather than quadratic. The different colors and shapes designate different mean competition strengths and the error bars denote one standard deviation above and below the average value. Each point is an average of 10 realizations. In all panels, there are $S = 30$ species and the average growth rate is $\mu_R = 1.5$. The columns designate different variabilities in growth rates, while the rows show different parameterizations of the self-regulation parameter $d_i$. In the constant row, $d_i = 1$ for all cases, while in the mean-adjusted row, $d_i = 1 + \mu_B/S^2$ so that varying the mean higher-order interaction strength changes all higher-order terms (including the cubic higher-order interactions). We simulated communities with a very large mean competition strength ($\mu_B = 20$ in the blue points) to more clearly illustrate the qualitative effect of competition in this model.

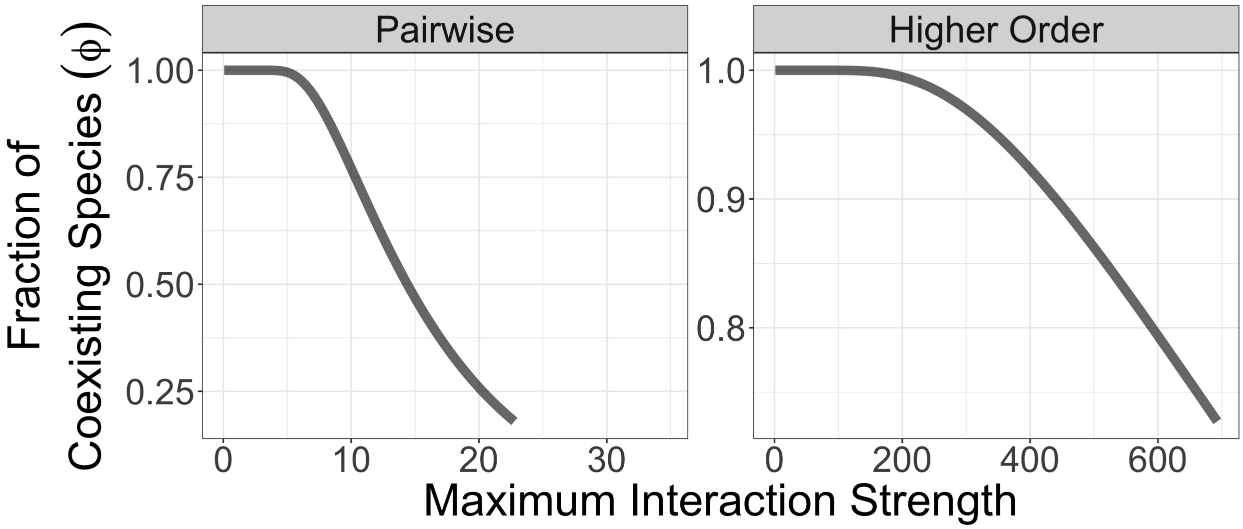

Figure N: **Effect of purely competitive interactions on species coexistence** We plot the predicted fraction of coexisting species for communities with only pairwise or only higher-order interactions when both of these interactions are constrained to be purely competitive. Because there is now a relationship between the mean and the variance of the underlying distributions, we now vary the maximum interaction strength, rather than varying the mean and variance independently. Parameters: $S = 20$, $\mu_R = 1.5$ and $\sigma_R = 0$.

#### References

[1] Mézard, M., Parisi, G. & Virasoro, M. A. SK Model: The Replica Solution without Replicas. *Europhysics Letters (EPL)* **1**, 77–82 (1986). URL https://doi.org/10.1209/0295-5075/1/2/006. Publisher: IOP Publishing.

[2] Bunin, G. Ecological communities with Lotka-Volterra dynamics. *Physical Review E* **95**, 042414 (2017). URL https://link.aps.org/doi/10.1103/PhysRevE.95.042414. Publisher: American Physical Society.

[3] Barbier, M. & Arnoldi, J.-F. The cavity method for community ecology. preprint, Ecology (2017). URL http://biorxiv.org/lookup/doi/10.1101/147728.

[4] Barbier, M., Arnoldi, J.-F., Bunin, G. & Loreau, M. Generic assembly patterns in complex ecological communities. *Proceedings of the National Academy of Sciences* **115**, 2156–2161 (2018). URL https://www.pnas.org/content/115/9/2156. Publisher: National Academy of Sciences Section: Biological Sciences.

[5] Wilson, W. G. *et al.* Biodiversity and species interactions: extending Lotka–Volterra community theory. *Ecology Letters* **6**, 944–952 (2003). URL https://onlinelibrary.wiley.com/doi/abs/10.1046/j.1461-0248.2003.00521.x. _eprint: https://onlinelibrary.wiley.com/doi/pdf/10.1046/j.1461-0248.2003.00521.x.

[6] Wilson, W. G. & Lundberg, P. Biodiversity and the Lotka–Volterra theory of species interactions: open systems and the distribution of logarithmic densities. *Proceedings of the Royal Society of London. Series B: Biological Sciences* **271**, 1977–1984 (2004). URL https://royalsocietypublishing.org/doi/10.1098/rspb.2004.2809. Publisher: Royal Society.

[7] AlAdwani, M. & Saavedra, S. Is the addition of higher-order interactions in ecological models increasing the understanding of ecological dynamics? *Mathematical Biosciences* **315**, 108222 (2019). URL http://www.sciencedirect.com/science/article/pii/S0025556418307247.

[8] Pettersson, S., Savage, V. M. & Nilsson Jacobi, M. Predicting collapse of complex ecological systems: quantifying the stability–complexity continuum. *Journal of The Royal Society Interface* **17**, 20190391 (2020). URL https://royalsocietypublishing.org/doi/full/10.1098/rsif.2019.0391. Publisher: Royal Society.

[9] Pettersson, S., Savage, V. M. & Jacobi, M. N. Stability of ecosystems enhanced by species-interaction constraints. *Physical Review E* **102**, 062405 (2020). URL https://link.aps.org/doi/10.1103/PhysRevE.102.062405.

[10] Bunin, G. Interaction patterns and diversity in assembled ecological communities. *arXiv:1607.04734 [cond-mat, physics:physics, q-bio]* (2016). URL http://arxiv.org/abs/1607.04734. ArXiv: 1607.04734.

[11] Barbier, M., de Mazancourt, C., Loreau, M. & Bunin, G. Fingerprints of High-Dimensional Coexistence in Complex Ecosystems. *Physical Review X* **11**, 011009 (2021). URL https://link.aps.org/doi/10.1103/PhysRevX.11.011009.
